# Genome-Scale Metabolic Model Accurately Predicts Fermentation of Glucose by *Chromochloris zofingiensis*

**DOI:** 10.1101/2021.06.22.449518

**Authors:** Michelle Meagher, Alex Metcalf, S. Alex Ramsey, Walter Prentice, Nanette R. Boyle

## Abstract

Algae have the potential to be sources of renewable fuels and chemicals. One particular strain, *Chromochloris zofingiensis*, is of interest due to the co-production of triacylglycerols (TAGs) and astaxanthin, a valuable nutraceutical. To aid in future engineering efforts, we have developed the first genome-scale metabolic model on *C. zofingiensis*, iChr1915. This model includes 1915 genes, 3413 metabolic reactions and 2652 metabolites. We performed detailed biomass composition analysis for three growth conditions: autotrophic, mixotrophic and heterotrophic and used this data to develop biomass formation equations for each growth condition. The completed model was then used to predict flux distributions for each growth condition; interestingly, for heterotrophic growth, the model predicts the excretion of fermentation products due to overflow metabolism. We confirmed this experimentally via metabolomics of spent medium and fermentation product assays. An *in silico* gene essentiality analysis was performed on this model, as well as a flux variability analysis to test the production capabilities of this organism. In this work, we present the first genome scale metabolic model of *C. zofingiensis* and demonstrate its use predicting metabolic activity in different growth conditions, setting up a foundation for future metabolic engineering studies in this organism.

## Introduction

Microalgae have the potential to serve as cellular factories, converting sunlight and carbon dioxide into a variety of valuable chemicals including lipids and fatty acids for fuel production. Rewiring metabolism for the overproduction of lipids or other metabolites is a difficult task which typically requires several changes in gene expression. Editing metabolic pathways while avoiding deleterious effects on the health of the cell requires extensive knowledge of the metabolic reaction network; specifically, how it functions and how it is regulated. One tool that can provide this critical information is a stoichiometric model of cellular metabolism. These models are relatively easy to generate and use and provide valuable insight into the metabolic network of a cell. Starting with a genome sequence, automated network annotation programs [1-3] can be used to create a high quality first draft model which enumerates all the metabolic reactions in the cell. After manual curation, the model can then be used to predict intracellular carbon flux distributions. Genome-scale metabolic models (GSMs) have been used successfully in a number of engineering efforts to design gene editing strategies in algae [4] and bacteria [5-11]. Therefore, the logical first step in utilizing microalgae as cell factories is to develop the computational models necessary to interrogate metabolism and model the impact of genetic changes.

*Chromochloris zofingiensis* has a fully sequenced genome [12] and has been identified as a lead candidate for the production of lipids for fuel production [13]. *C. zofingiensis* has been reported to accumulate up to 40% of its cell dry weight as triacylgylcerols (TAGs) [14] and it can also accumulate high levels of astaxanthin [15, 16], a naturally occurring pigment with powerful antioxidant properties. Due to its stereospecificity and antioxidant capabilities, naturally produced astaxanthin is a high-value nutraceutical with an estimated value of $7,000/kg [17]. The co-production of TAGs and a high-value product like astaxanthin make the economic feasibility of biofuels production from algae much more attainable. Numerous studies have explored growth conditions to maximize the production of lipids and astaxanthin in the cell. Stress conditions have been shown to increase expression of astaxanthin biosynthesis related genes in *C. zofingiensis* [18] and indeed growth in stress conditions (high light, nitrogen deprivation, high salinity) results in elevated levels of astaxanthin [19-22]. Stress conditions have also been associated with higher lipid content [19], but this comes at the cost of a lower biomass productivity [23]. *C. zofingiensis* is capable of growing heterotrophically on various sugars, and heterotrophic growth is also associated with elevated levels of astaxanthin in the cell [24-26]. Recent studies have shed light on the metabolic activities of *C. zofingiensis* when grown on glucose, by analyzing both the transcriptome response and physical changes to the cell that occur when glucose is introduced. It was found that *C. zofingiensis* is capable of a unique and reversible metabolic shift from autotrophic to purely heterotrophic growth when provided glucose [27]. This trophic switch is characterized by the downregulation of chlorophyll production, loss of photosynthetic machinery and alterations in the thylakoid membrane structure, accompanied by the accumulation of energy storage compounds such as lipids and starch [27]. The downregulation of chlorophyll synthesis leads to a decreased concentration of primary photosynthetic pigments within the cell, even when grown under continuous illumination. Unlike nitrogen deprivation induced accumulation of lipids and astaxanthin, which suffers from decreased biomass productivity [23], this reversible glucose initiated trophic switch results in an increase in biomass productivity [27]. In this work, we present the first genome-scale model of *C. zofingiensis*, and its use to predict growth and carbon flux distributions in three different growth conditions: autotrophic, heterotrophic and mixotrophic. We have performed a detailed biochemical analysis of cellular biomass composition for each of these growth conditions to formulate accurate biomass formation equations. The results from these simulations predicted the formation of fermentation products in cultures grown on glucose, a previously uncharacterized behavior in this alga that has since been confirmed through two different methods of analysis.

## Results

### Network Reconstruction and Curation

The initial model draft was constructed using RAPS, an automated algorithm for building high quality genome scale metabolic models of new algal species. In this case, the donor models used to construct the model of *C. zofingiensis* were *Chlamydomonas reinhardtii* [28] and *Nannochloropsis gaditana* [29]. The final curated model, iChr1915, consists of 1915 genes and 3413 reactions (including 490 transport reactions). Figure 1 summarizes model details, including reactions shared with donor models and compartmentalization of reactions and metabolites. Full lists of metabolites and reactions included in the model are included as supplemental files 2 and 3, respectively. The draft model created by RAPS was manually curated to ensure accuracy. Tables S1 and S2 in the Supplemental Information (SI) provide a list of reactions that were manually added or removed from the RAPS output model during manual curation. Less than 2% of the model required manual curation; most notably, astaxanthin synthesis was added manually as neither donor model included these steps. After manual curation, the model had a MEMOTE score of 84%, indicating a consistent model with good genomic support and annotations [30]. More detailed information on the MEMOTE score of this genome scale model can be found in supplemental file 10.

**Figure 1.**
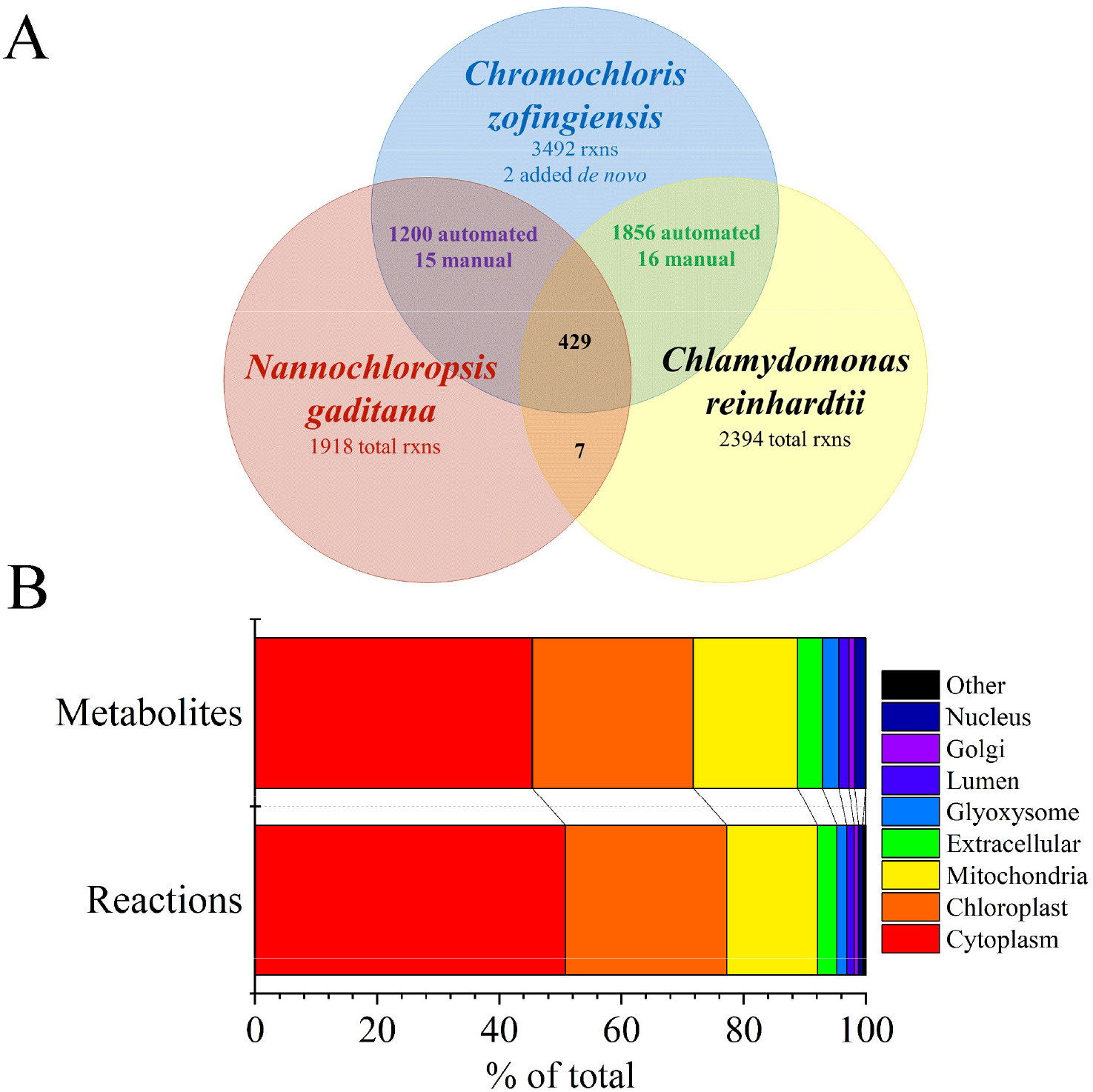
Details of the *Chromochloris zofingiensis* genome-scale metabolic model. RAPS [1] was used to construct the genome scale metabolic model from two donor models. A) Contribution of reactions from donor models to the initial genome-scale metabolic model, before curation. The finalized model contained 3413 reactions, 2652 metabolites, and 8 cellular compartments. B) Localization of reactions and metabolites. Reactions for biomass generation and spectral decomposition of light sources are included in the “other” category.

### Biomass Composition and the Biomass Objective Function

We measured the biomass composition of cells in mid-exponential growth in continuous light (see Figure 2) for all growth conditions. This data, along with some key assumptions based on literature data on lipid and pigment compositions [28] were used to construct detailed biomass formation equations for each growth condition. It was assumed that the general profile of lipids and carotenoid pigments resembled that of *Chlamydomonas reinhardtii*, but with the inclusion of astaxanthin. A table of the full biomass formation equation for each growth condition is provided as Table S4 in the supplemental information.

**Figure 2.**
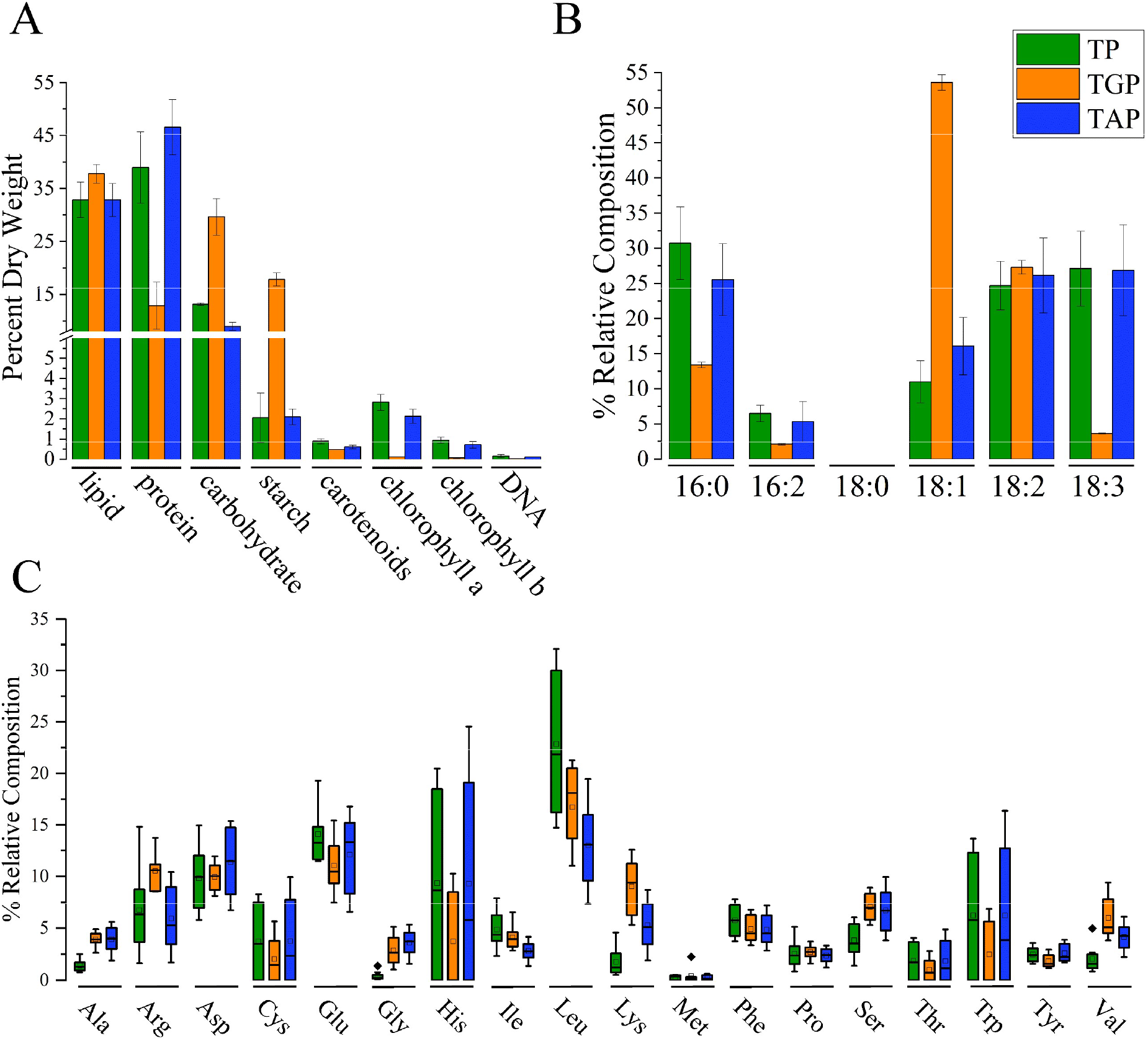
Average biomass composition measurements of *Chromochloris zofingiensis* in three different media conditions: minimal media (TP), mixotrophic growth with acetate (TAP) and heterotrophic growth with glucose (TGP). A) Biomass composition in terms of major macromolecules, B) Fatty acid composition and C) amino acid composition of hydrolyzed proteins. Error bars represent standard deviation of n = 3 samples.

### Experimentally measured fluxes and rates used to constrain the model

To simulate *in vivo* growth conditions accurately, we measured reduced carbon uptake rates as well as the growth rate for each condition experimentally (Table 1) and used these as constraints for the pFBA simulation. Experimental data used to define these constraints is presented in the supplemental information as Figures S2, and a full list of pFBA constraints is given in Table S3. As expected, growth in the presence of a reduced carbon source increases the growth rate; the doubling time in the presence of glucose is approximately 16.5 hours compared to 25.3 hours in acetate and 26.3 hours for photoautotrophic growth.

**Table 1.** Experimentally measured fluxes and rates used to constrain the metabolic model.

| Model Constraints | Media Formulation |  |  |
| --- | --- | --- | --- |
|  | TP | TAP | TGP |
| Growth Rate ($\text{h}^{-1}$) | $0.0264 \pm 0.0004$ | $0.0274 \pm 0.0025$ | $0.042 \pm 0.0089$ |
| Acetate ($\text{mmol/gDW-hr}$) | | $3.22 \pm 0.78$ | |
| Glucose ($\text{mmol/gDW-hr}$) | | | $2.41 \pm 0.11$ |

### Flux Distributions and Predicted Exchanges

We simulated growth in three conditions: photoautotrophic (TP medium), mixotrophic with acetate (TAP medium) and heterotrophic with glucose (TGP medium). Although continuous light was used in all data collection experiments, it has been shown that *Chromochloris zofingiensis* degrades its photosynthetic apparatus in the presence of glucose, and this loss of photosynthetic activity is further evidenced by measured F_v_/F_m_ values decreasing from 0.66 to 0.01 upon the addition of glucose [27]. Therefore, regardless of light conditions, cultures provisioned with glucose will grow heterotrophically. Interestingly, this is reinforced by parsimonious flux balance analysis (pFBA) predictions; when growth in TGP medium is simulated, the pFBA solution predicts no absorption of light despite the model allowing it (Figure 5. **Predicted fluxes for heterotrophic growth of *C. zofingiensis* on glucose**. The thickness of the arrows indicates the amount of flux through the given reaction on a linear scale as shown in the legend in the lower right. The model is compartmentalized, and this is reflected in the figure: cytosol is shown in white, plastid in green and mitochondria in orange. Arrows in gray indicate viable reactions that currently carry negligible flux. This figure was generated automatically by the mapping tool associated with RAPS [1].). Simulations for growth on glucose and acetate also predict an array of fermentation products, including acetate, lactate, succinate, ethanol, and formate. The flux map for autotrophic growth (Figure 3) shows the majority of carbon flux is directed through the Calvin Benson Bassham (CBB) cycle in the chloroplast, as would be expected. There is also significant flux through malate valves to shuttle ATP and NAD(P)H into the different cellular compartments [31]. In mixotrophic growth (Figure 4), far less flux is seen through the CBB cycle and instead triose phosphates are transported into the cytosol and are shuttled back and forth between the mitochondria to move energy and reducing equivalents around the cell. Acetate taken up by the cell is transported into the mitochondria and converted into acetyl-coA, before being converted into biomass and some fermentation products. This indicates the cell is likely consuming acetate as a rate greater than required for the measured growth, and therefore has excess energy available; additionally, this energetic excess is magnified when grown on glucose. In heterotrophic growth (Figure 5), the model correctly predicts no photosynthetic light harvesting (without constraints placed on the model to limit/eliminate it) [27]. Although glycolytic enzymes are present in both the plastid and cytosol, the model predicts that glucose is catabolized in the plastid (Figure 5); we hypothesize this is due to the need for ATP and NADH in the plastid. The model also predicts that growth on glucose induces excretion of several overflow metabolites, such as acetate, lactate, succinate and ethanol (Table 2), suggesting that the cell is uptaking more reduced carbon than might be expected if the cell was fully utilizing the TCA cycle. Each of these flux maps indicate that there are significant differences between the growth conditions modeled and that these manifest as differences in biomass composition (Figure 2).

**Table 2.** Predicted Uptake and Excretion Fluxes. Negative values indicate uptake, while positive values indicate excretion.

| **Predicted Exchanges**<br>**(mmol/gDW-hr)** | **TP** | **TAP** | **TGP** |
| --- | --- | --- | --- |
| Oxygen | 0 | -2.34 | -0.237 |
| Carbon Dioxide | -1.17 | 2.97 | 3.87 |
| Water | 0 | 2.98 | 0 |
| Acetate | 0 | -3.22 | 1.29 |
| Lactate | 0 | 0.410 | 0.0085 |
| Succinate | 0 | 0.0122 | 0.332 |
| Ethanol | 0 | 0.395 | 2.39 |
| Formate | 0 | 0.0011 | 0.0015 |
| Photons | -13.9 | -0.0003 | -0.0004 |

**Figure 3.**
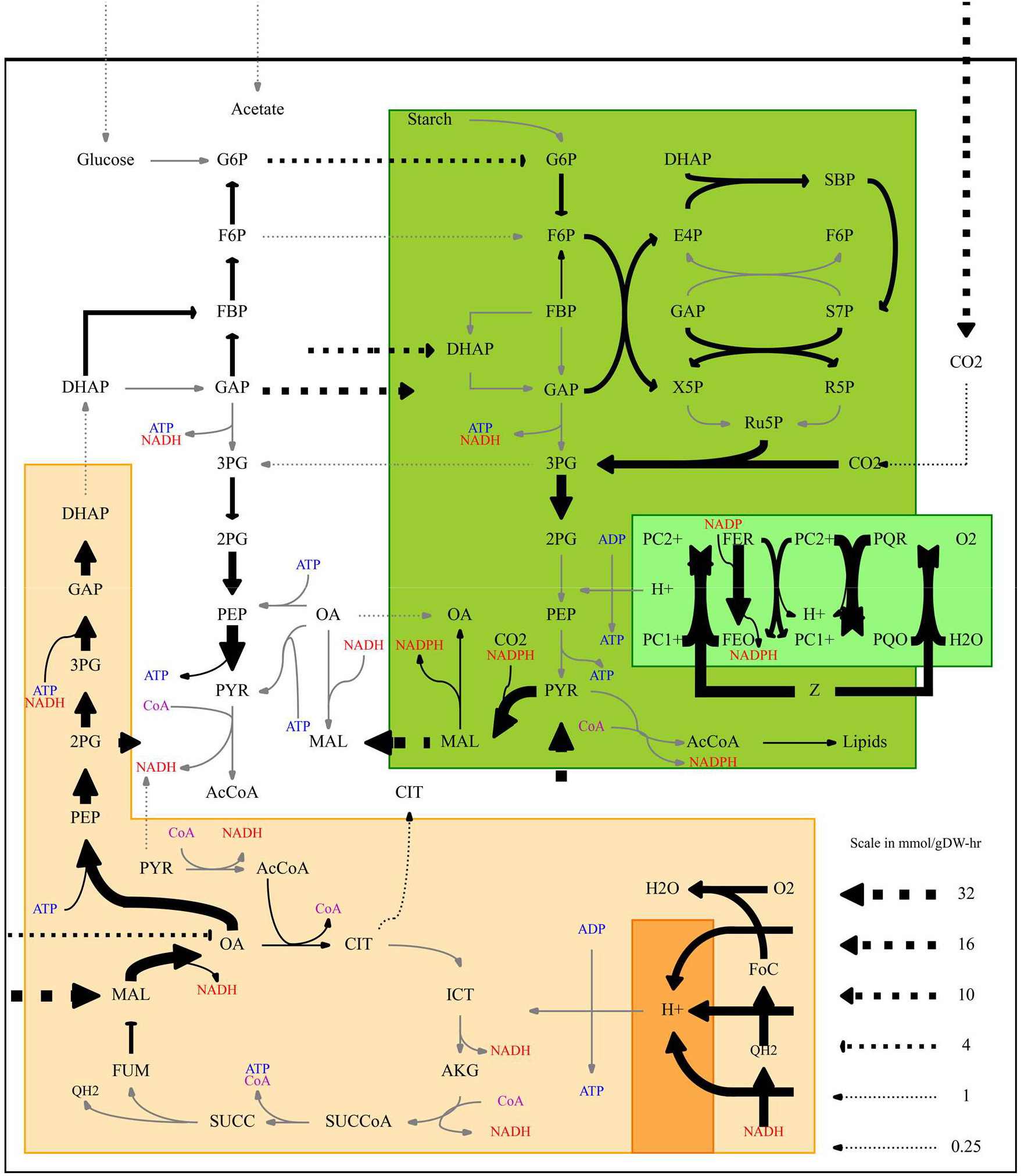
Predicted fluxes for photoautotrophic growth of *C. zofingiensis*. The thickness of the arrows indicates the amount of flux through the given reaction on a linear scale as shown in the legend in the lower right. The model is compartmentalized, and this is reflected in the figure: cytosol is shown in white, plastid in green and mitochondria in orange. Arrows in gray indicate viable reactions that currently carry negligible flux. This figure was generated automatically by the mapping tool associated with RAPS [1].

**Figure 4.**
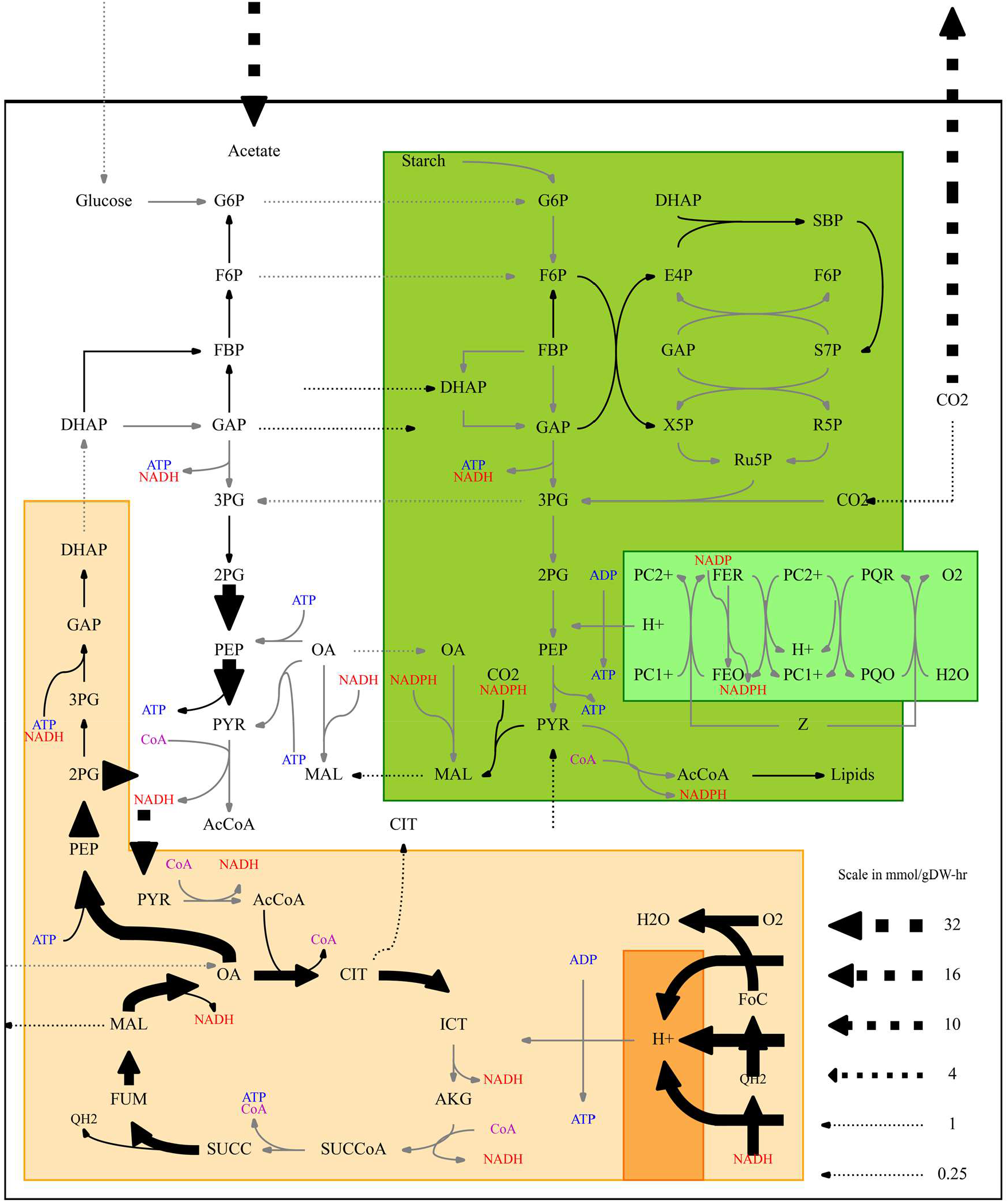
Predicted fluxes for mixotrophic growth of *C. zofingiensis* on acetate. The thickness of the arrows indicates the amount of flux through the given reaction on a linear scale as shown in the legend in the lower right. The model is compartmentalized, and this is reflected in the figure: cytosol is shown in white, plastid in green and mitochondria in orange. Arrows in gray indicate viable reactions that currently carry negligible flux. This figure was generated automatically by the mapping tool associated with RAPS [1].

**Figure 5.**
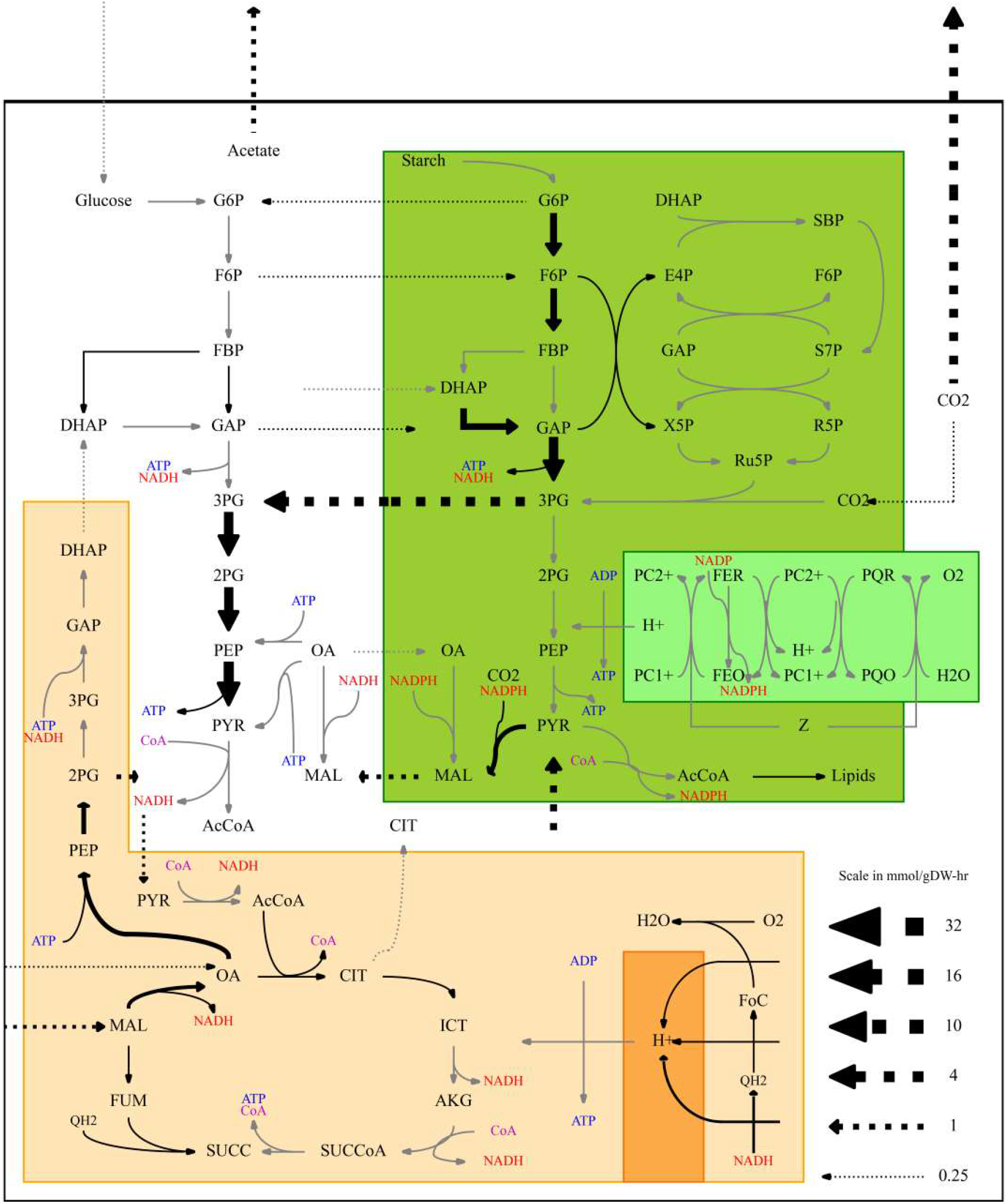
Predicted fluxes for heterotrophic growth of *C. zofingiensis* on glucose. The thickness of the arrows indicates the amount of flux through the given reaction on a linear scale as shown in the legend in the lower right. The model is compartmentalized, and this is reflected in the figure: cytosol is shown in white, plastid in green and mitochondria in orange. Arrows in gray indicate viable reactions that currently carry negligible flux. This figure was generated automatically by the mapping tool associated with RAPS [1].

### Metabolomics Analysis

To further investigate the presence of fermentation product excretion, we performed untargeted metabolomics of spent media from cultures grown in TAP and TGP, sampled in late exponential phase, to identify metabolites excreted during growth. Results of this analysis are presented in Figure 6, showing the peak areas for select metabolites that are found at much higher concentrations in TGP than in TAP media. The full dataset is provided as Supplemental File 4. The most abundant metabolite found in the media was lactate, followed by the amino acid leucine.

**Figure 6.**
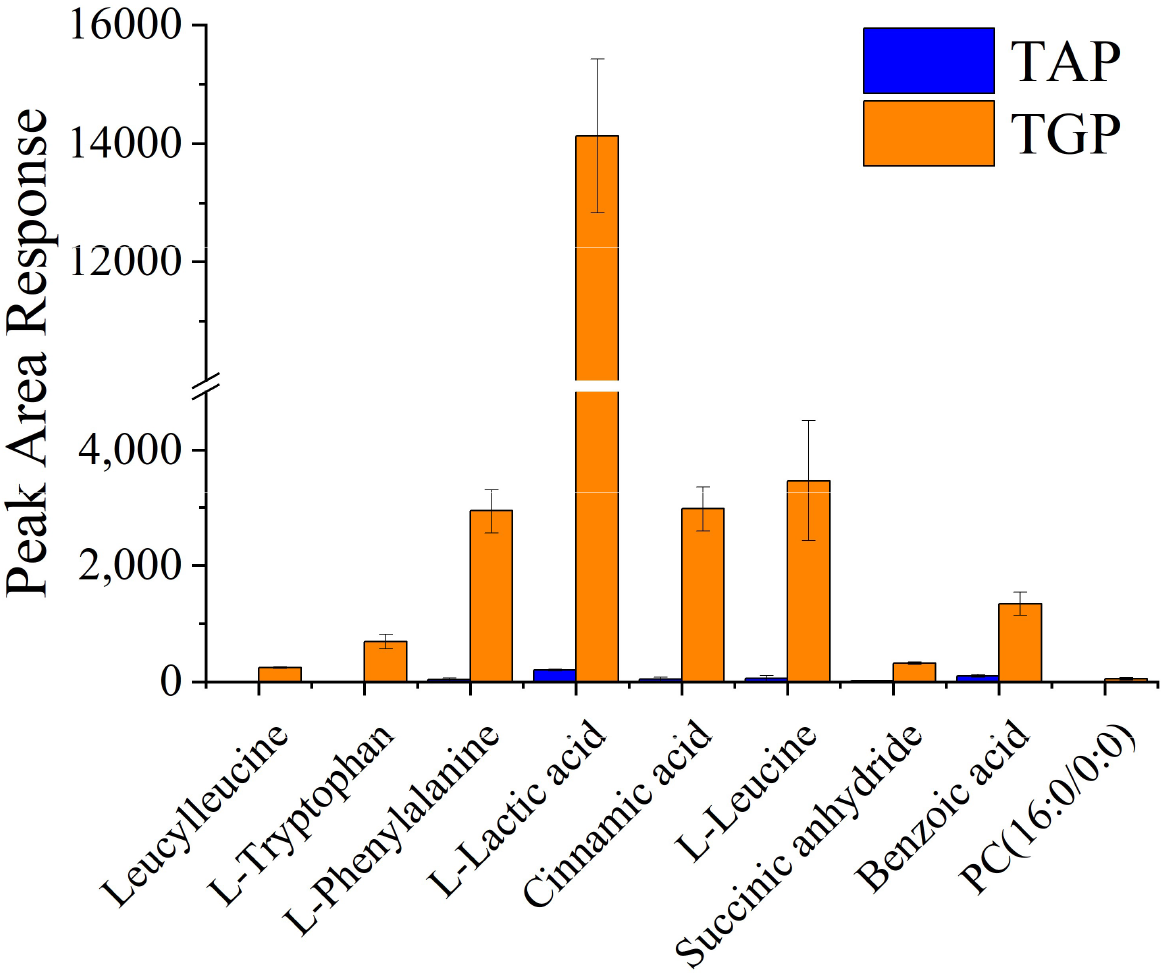
Peak areas for selected metabolites measured via ultraperformance liquid chromatography mass spectrometry in electrospray ionization (ESI) negative mode in TAP and TGP spent media. Extracellular media in TGP cultures was found to contain a mixture of organic acids, amino acids, and energy storage compounds in significantly higher amounts than that of TAP cultures. Lactic acid appears as the predominate metabolite being exported.

Our metabolomics analysis was not quantitative, so we followed up the metabolomics analysis with an analysis of filtered media samples conducted on a YSI biochemistry analyzer. This data is presented in supplemental Figure S3. The lactate excretion rate observed was 0.0713 ± 0.004 mmol/gcdw/h; this is more than was predicted by the pFBA output of the model, but it is still well within thermodynamic bounds. Lactate was not detected in TAP spent media samples.

### Essential Gene Analysis

An *in silico* gene essentiality analysis was performed on the genome-scale metabolic model of *Chromochloris zofingiensis* for growth in continuous light. The results of this analysis indicated that out of the 1914 genes included in the model, only 82 were essential for growth in both the autotrophic and mixotrophic cases (essential genes reduce growth to 10% or less when knocked out) while 93 genes were essential for growth in the heterotrophic case. Table 4 shows the summarized results of this analysis; the full list of essential genes for each growth condition is provided as Table S5 in the supplemental file.

Unsurprisingly, both autotrophic and mixotrophic growth both require genes associated with the light harvesting apparatus, while heterotrophic growth requires the genes associated with mitochondrial electron transport chain. Other than the examples listed above, there were 81 genes essential for growth in all growth conditions. This list included transketolase genes associated with reactions in the pentose phosphate pathway and the gene for Mg^2+^ transporters. Other genes essential for all growth conditions were involved in the production of biomass. These genes can be grouped into amino acid and protein synthesis (35), nucleic acid synthesis (18), lipid synthesis (11), pigment synthesis (10), and energy molecule synthesis (5).

### Flux Variability Analysis

As heterotrophic growth triggers accumulation of lipids and astaxanthin, examining the metabolic constraints of that may yield insights into possible targets of genetic manipulations. We used the model to evaluate flux variability in different growth conditions that may lead to increases in production of lipids or astaxanthin: heterotrophically. We simulated five different cases – TGP, TGP with lipid export at both full biomass production and no biomass production, and TGP with astaxanthin export at both full biomass production and no biomass production (Figure 7**Error! Reference source not found**.). The uppermost bar is the normal TGP biomass production case, where the growth rate is fixed to experimental values and there is no additional production of any desired products. The next two bars show the values for maximal production of triolein (a lipid composed of only oleic acid tails, chosen because of the high fraction of oleic acid produced by TGP) and astaxanthin, respectively, without any required biomass production. The last two represent the maximal production of triolein and astaxanthin with the TGP fixed growth rate. The increase in flux through lipid and carotenoid pathways is unsurprising, as these pathways represent the sole mechanisms for moving more carbon into the desired products. Moreover, the cell’s use of fermentation is clearly visible in these FVA results, as there is a wide range of potential flux values for the nominal TGP biomass production case, indicating that the cell is not limited by energy. A full set of FVA data for all reactions is available in supplemental files 5-9.

**Figure 7.**
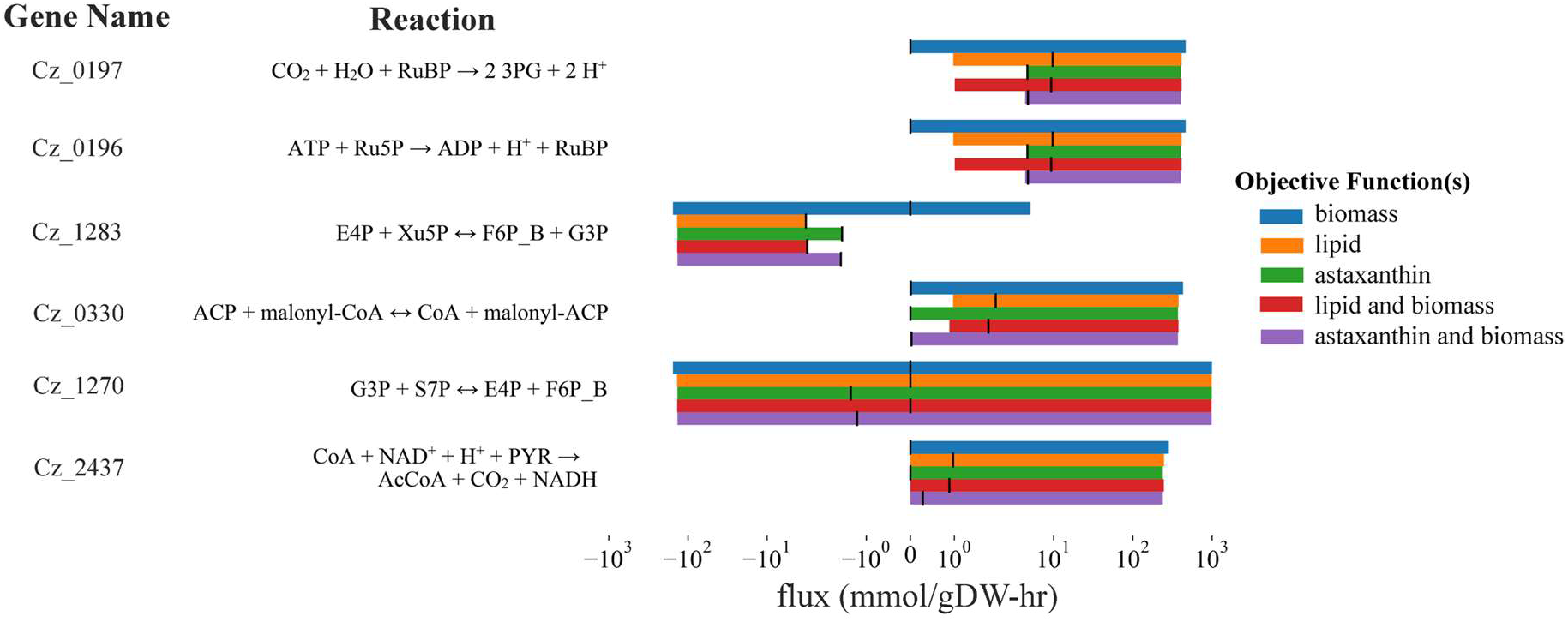
Example flux distributions and variabilities through several reactions in *C. zofingiensis* under TGP conditions. The “Biomass” case is the regular fixed TGP production case, while the “Lipid” and “Astaxanthin” cases represent the maximal production of triolein and astaxanthin, respectively, both with and without biomass growth. The black lines are the pFBA solutions for each case, while the bars are the FVA ranges. Unsurprisingly, astaxanthin production clearly increases flux through parts of the pentose phosphate pathway, while lipid production relies heavily on shuttling carbon through malonyl derivatives.

## Discussion

### Creation of the Genome Scale Metabolic Model

The creation of a genome-scale model is often an iterative process; as more is learned about the organism of interest and genome annotation improves, the model improves as well. Typically, this means that each subsequent use of the model results in a more complete and/or more accurate model based on new discoveries. For the development of the *Chromochloris* model, we used RAPS RAPS, an automated algorithm specifically designed for the creation of algal GSMs, to create the first draft network in less than an hour. The first draft network was then subject to manual curation to remove duplicate or erroneous reactions, add missing reactions and fill gaps in pathways. The manual curation required is dependent on the quality of the donor models used in RAPS, and their similarity to the organism of interest. Presently, there are a limited number of genome scale metabolic models of algae to select from for use as RAPS donor models, so manual curation of the RAPS output was necessary. Manually added reactions included the two reactions required for astaxanthin synthesis from zeaxanthin (the production of adonixanthin from zeaxanthin and the production astaxanthin from adonixanthin), as those reactions are not included in the RAPS source models.

### Biomass Composition

Autotrophic and mixotrophic cultures supplemented with acetate were observed to have similar growth rates and phenotypes. This is also confirmed with the similarity in biomass composition. The biomass composition of heterotrophic cultures were significantly different than both autotrophic and mixotrophic cultures in biomass composition, growth characteristics, and physical appearance. This suggests that significant metabolic shifts occur in *C. zofingiensis* when grown heterotrophically. In terms of macromolecular composition, growth on glucose leads to higher lipid and starch content, and lower protein content (see Figure 2A). This tendency for *C. zofingiensis* to accumulate carbon storage compounds when grown in the presence of glucose is well documented in the literature [25, 27, 32-34]. Our average experimental value of lipid content in heterotrophic cultures of *C. zofingiensis*, expressed as percent dry weight, was 37.8% ± 1.7. This lies in agreement with values reported in the literature ranging from 30 - 52% lipid content on a cell dry weight basis, depending on the concentration of sugar provided in the medium [25, 32-34]. Heterotrophic cultures grown on glucose were found to have lower protein content than cultures grown in other media conditions. It has been observed that *C. zofingiensis* cultures grown on glucose experience a loss of photosynthetic machinery [27]. Additionally, the gene essentiality analysis conducted on *C. zofingiensis* presented in this work indicates that the gene encoding RubisCO is not essential for growth on glucose. In *Chlamydomonas reinhardtii*, the photosynthetic light harvesting apparatus and RubisCO combined account for 13.9% of total cellular protein content [35], so it is possible that the lower protein content observed in cultures grown on glucose is the result of the loss of photosynthetic proteins in these cultures. We also measured an increase in the oleic acid content of the lipid fraction (see Figure 2B), which agrees with previous studies [25, 32]. A study by Liu et al. reported that the introduction of glucose in *C. zofingiensis* cultures led to a dramatic increase in the expression of the genes encoding stearoyl-acyl carrier protein desaturase (SAD) and the BC subunit of acetyl-CoA carboxylase (ACCase) [36]. ACCase catalyzes the first step in fatty acid synthesis, and SAD catalyzes the conversion of stearic acid to oleic acid, therefore, this change in gene expression is likely the reason for the high oleic acid content observed.

### Flux Maps

FBA simulations were performed to quantify changes in flux distributions resulting from growth on different media formulations. Autotrophic growth has high flux through photosynthetic light reactions, carbon fixation, and the pentose phosphate pathway (Figure 3). In autotrophic conditions, the flux distribution is very similar to that calculated for *Chlamydomonas reinhardtii* [37]. However, flux distributions calculated for mixotrophic growth on acetate differ from that in *C. reinhardtii* [37], with zero flux through the TCA cycle, and little carbon fixation activity (Figure 4). The acetate that is taken up by the cell is directed to both lipid synthesis and energy production. *Chromochloris zofingiensis* is not capable of mixotrophic growth on glucose, due to the degradation of the photosynthetic apparatus upon the introduction of glucose to the cultures [27]. The flux maps for heterotrophic growth show a high flux through glycolysis and have almost no flux through the pentose phosphate pathway (Figure 5); this agrees with another metabolic model of Eukaryotic algae, *Chlorella vulgaris*, which had no flux through the oxidative phase of the pentose phosphate pathway under heterotrophic conditions [38]. Interestingly, simulations for cultures grown mixotrophically on acetate and heterotrophically on glucose predicted the formation of organic acid fermentation products. A large portion of the glucose flux (61%) is excreted as fermentation products instead of being utilized by the TCA cycle, and similar behavior is predicted in mixotrophic cultures. Flux balance analysis [39] and metabolic flux analysis [40] on heterotrophically grown algae support the notion that the full TCA cycle may not be used during heterotrophic growth. [39, 40]. The model is not limited in anyway by oxygen uptake, so this is not obligatorily anaerobic fermentation – given the large amount of protein required for the TCA cycle, it is likely that the cell ferments glucose as a way to reduce the amount of enzyme required for the cell to produce energy.

Excess production capabilities were evaluated using flux variability analysis in Table 4. There are several critical points raised by this table. First, the presence of fermentation products in the “Measured Growth” result, and the corresponding lack of such in the “Maximum Growth” case. While ethanol is not the only fermentation product produced in this case, it is included as an example to highlight this switch. The cell’s fermentation is also predicted by the extremely low oxygen uptake in the “Measured Growth” model and the corresponding switch to high oxygen consumption in the “Maximum Growth” model. Fixed growth produces fermentation because the relatively low energy needs of the cell can be adequately met by the simple fermentative pathways, and the pFBA formulation of the model optimization will always produce the simplest (lowest total flux) solution.

This large energy excess appears to come from the cell’s preference for fermentation. Table 4. **Comparison of measured and maximum growth exchanges** compares the growth of the cell in two cases. Both are simulated in TGP, and therefore have identical rates of glucose uptake, but the “Maximum Growth” case does not have growth fixed to a specific value. Instead, it is unbounded and optimized for. This case represents the cell’s production potential when glucose can be utilized as efficiently as possible. The striking changes in the oxygen, carbon dioxide, and ethanol exchanges reinforce that the cell resides in a majority-fermentation mode under high glucose conditions. This large energy excess appears to come from the cell’s preference for fermentation. Table 4 compares the growth of the cell in two cases: both are simulated in TGP, and therefore have identical rates of glucose uptake, but the simulated maximum growth case allows the model to freely adjust growth rate. This case represents the cell’s biomass production potential when glucose can be utilized as efficiently as possible. The striking changes in the oxygen and carbon dioxide exchanges reinforce that the cell resides in a majority-fermentation mode under high glucose conditions – the fermentation products listed in Table 2 all vanish under maximal growth, as the cell can efficiently extract more energy by oxidizing the carbon into carbon dioxide, via the TCA cycle. These results present two conclusions: First, glucose transporters are not limiting the growth rate of the cell, as otherwise the rate of glucose consumption would be low enough to require non-fermentative energy generation. Second, something is limiting the cell’s ability to respire and efficiently utilize glucose. It may be valuable to conduct studies on oxygen levels in the culture, to determine if there is sufficient oxygen available to respire, as well as experiments in cultures with low glucose, to determine the cell’s transition point between fermentation and respiration.

### Network Evaluation and Robustness

An *in silico* gene essentiality analysis was performed to assess metabolic network robustness in *C. zofingiensis*. This was done by performing a series of continuous light single gene knockouts, simulating cell growth under these gene deletions, and finally comparing predicted growth rates to that of the wild type cells. Gene deletions which resulted in growth rates of >80%, 10-80%, and <10% of the wild type simulations enabled the genes to be classified as non-essential, beneficial, and essential. The genes encoding Ribulose-1,5-bisphosphate carboxylase-oxygenase (RuBisCO) was categorized as essential for growth in autotrophic and mixotrophic cases, but not for the heterotrophic case. In the case of a RuBisCO knockout in mixotrophic growth, the maximum growth rate that can be achieved is only 41% of the wild type growth rate, even though acetate uptake made up roughly two-thirds of the total wild-type carbon uptake. 12 genes were found to be essential in heterotrophic growth, but not in the autotrophic or mixotrophic cases, and all of these related to the ATP synthase complex V in the mitochondria. The high number of non-essential genes in each growth condition (see Table 3) points towards a highly robust metabolic network. This analysis was modeled after an *in silico* gene essentiality analysis performed by Broddrick et.al. [39]. In this study, the initial computational evaluation for the number of essential genes in *Synechococcus elongatus* was 134, which was later increased based on *in vivo* data collected [39]. It is likely that computational evaluations underestimate the number of essential genes for growth for several reasons [39], nonetheless, by comparing this data set to *in vivo* data on gene essentiality important knowledge gaps in the *C. zofingiensis* metabolic network may be identified.

**Table 3.** Results of the gene essentiality analysis for all three growth conditions. Genes which, when deleted, result in a simulated growth rate of >80%, 80-10%, and <10% of the wild type are categorized as non-essential, beneficial, and essential, respectively.

|  | **Autotrophic** | **Mixotrophic** | **Heterotrophic** |
| --- | --- | --- | --- |
| Essential | 82 | 82 | 93 |
| Beneficial | 14 | 16 | 15 |
| Non-essential | 1818 | 1818 | 1806 |

**Table 4.** Comparison of measured and maximum growth exchanges.

| Simulation | Growth Rate | Glucose Exchange | CO<sub>2</sub> Exchange | O<sub>2</sub> Exchange | Ethanol Exchange |
| --- | --- | --- | --- | --- | --- |
|  | (1/hr) | (mmol/gDW-hr) |  |  |  |
| experimentally determined growth rate and glucose uptake | 0.039 | -2.91 | 2.49 | -0.06 | 2.22 |
| simulated maximum growth rate with experimentally determined glucose uptake | 0.230 | -2.91 | 6.16 | -4.05 | 0 |

### Metabolomics analysis

The excretion of lactate to the medium in heterotrophic cultures was predicted by the model and validated by our metabolomics analysis. This has not been reported in the literature previously but is not completely unheard of. Algae in the *Chlamydomonas, Chlorogonium*, and *Chlorella* genera have been shown to be capable of fermentation [41], but this behavior is most often observed in anoxic conditions. In yeast, with high enough glucose concentrations, cells have been reported to preferentially ferment glucose even in the presence of oxygen [42, 43]. It has been postulated that this fermentation activity primarily occurs to re-generate NAD^+^/NADP^+^ used in glycolysis and balance the redox state of the cell [41]. To investigate the production of fermentation products, we performed an untargeted metabolomics experiment, analyzing spent media from heterotrophic cultures (TGP media) and mixotrophic cultures (TAP media). Lactate was identified in the extracellular TGP media in significant quantities, seemingly verifying the model prediction of excreted fermentation products. Interestingly, while this model prediction was validated for heterotrophic TGP cultures, no lactate was detected in TAP spent media samples. The presence of lactate was also observed in a separate analysis, with lactate concentrations in the extracellular TGP media reaching values of 1.96 mM, while lactate concentrations in TAP extracellular media remained below detection limits. Amino acids were also found to be present in the extracellular media, in far higher concentrations in heterotrophic cultures than mixotrophic cultures. It is also possible that observed extracellular metabolites are the result of cell lysis induced in lipid accumulating cultures. However, two metabolites that would be expected to increase in concentration from cell lysis, deoxyribose sugars and oleic acid, were found in lower levels in spent TGP media than TAP media. The high abundance of metabolites displayed, and lack of other intracellular metabolites is indicative that there is export of these compounds happening.

## Conclusion

This work presents iChr1915, the first genome scale metabolic model for the emerging model species *C. zofingiensis* and demonstrates its usefulness in making predictions of phenotype under different growth conditions. This organism is of high promise for biofuel production when grown heterotrophically, with both high lipid productivities and a fatty acid profile acceptable for fuel production [25]. We also present a detailed analysis of the biomass composition in *C. zofingiensis* in three different growth conditions, and an assessment of the metabolic network flexibility. Analysis and testing of the model indicated that *C. zofingiensis* preferentially ferments in the presence of glucose – and experimental evidence later confirmed this prediction. This successful validation means the model is useful for predicting cellular phenotypes under varying nutrient conditions. This model will be extremely useful in further understanding the metabolic implications of *C. zofingiensis*’s propensity to accumulate astaxanthin, and generating strategies to improve strain production.

## Materials and Methods

### Culture Growth Conditions

*Chromochloris zofingiensis* cultures provided by Professor Niyogi’s lab at UC Berkeley were grown in 500 mL baffled flasks in culture volumes of 200 mL. Cultures were maintained under constant light, with an average intensity of 90 μmol/ m^2^·s, a temperature of 25°C, and an agitation speed of 180 rpm. *C. zofingiensis* cultures were grown in three different media conditions, Tris-Phosphate (TP) minimal medium (adapted from [44]), TP supplemented with 25 g/L glucose (TGP), and TAP medium [44], each at a pH of 7. Specific growth rates for each set of culture conditions were measured in triplicate.

### Model Construction and Curation

An automated tool, Rapid Annotation of Photosynthetic Systems (RAPS) [1], was used to build the initial draft metabolic network reconstruction. This tool compares the cell’s predicted proteins to known proteins in other previously published models. The donor models used were eukaryotic microalgae whose genomes had been previously annotated, as detailed by Metcalf [1].

The major steps in model curation were the manual adjustments to the RAPS reactions, and the addition of the astaxanthin production reactions. There were several manual adjustments, as shown in Table S1 and Table S2. These modifications were generally performed to remove thermodynamically infeasible loops or redundant lumnial or intermitochondrial space reactions. Once RAPS had produced the draft model, the reactions for astaxanthin production were added.

### Biomass Composition Measurements

Culture samples were collected during mid exponential growth phase (See supplementary information, figure S1) for dry weight determination and biomass composition analysis. Cell dry weight was determined by vacuum filtration of 15 mL culture samples through a pre-dried, 1.6 μm pore size glass microfiber filter. After filtration, cells were rinsed with TP medium and dried in an 80°C oven overnight. For biomass composition analysis, culture volumes corresponding to approximately 5 mg cell dry weight were aliquoted and spun down to collect a cell pellet. Cell pellets were then ground with 5 mm diameter stainless steel balls in a Retsch cryomill to disrupt the tough cell wall observed in *C. zofingiensis*.

For analysis of protein content, the pellet from chlorophyll extraction was solubilized by suspending in 0.2 N NaOH and heating to boiling for 1 hr. A 1:5 dilution of this protein solution was quantified using a Pierce BCA protein assay kit (Thermo scientific). For analysis of the relative composition of amino acids, proteins were hydrolyzed, and the resulting mixture of amino acids was converted to their tert-butyldimethylsilyl derivatives as described in work by Antoniewicz et. al. [45]. The time allowed for the derivatization reaction was extended as described by Mawhinney et.al. [46] to overcome steric hindrance involved in the addition of the tert-butyldimethylsilyl group on histidine, tryptophan, and threonine. After derivatization, the mixture was run on the GC/MS with a quadrupole FID detector. Due to the significant decrease in response for histidine after 6 h noted by Mawhinney et.al. [46], it was ensured that all samples were run within 6 h of the completion of the derivatization reaction. A combination of amino acid standard solution (AAS18) from Sigma, and a prepared solution of 5 μM Tryptophan and Cysteine in 0.1 M HCl were used to generate standard curves for quantification. GC/MS conditions were as described by Antoniewicz et. al. [45].

Lipid content was determined by gravimetric analysis, using a chloroform: methanol extraction of ground cells as described in Breuer et. al. [47]. Fatty Acid composition was determined by converting fatty acids present in lysed cell material to their methyl ester derivatives (FAMEs) and analyzing the FAMEs on GC/MS. The procedure used for this analysis was outlined by Christie et.al. [48]. The proportions of lipid existing as triacylglycerides, phospholipids, and sulfolipids was assumed to be relatively consistent with those values measured in *Chlamydomonas reinhardtii*.

For starch and total carbohydrate analysis, pigments were extracted from a ground cell pellet with a 95% ethanol solution, which was then washed and re-suspended in 100 mM sodium acetate (pH 4.5). Samples were autoclaved to transform particulate starch to colloidal starches. For starch content determination, 100 μL of an 80 unit/mL solution of amyloglucosidase (Sigma Aldrich, S9144) was added to each autoclaved sample. Samples were then incubated at 55°C overnight to digest the starch into glucose, which was then quantified using a glucose assay (Sigma Aldrich, G3293). For determination of total carbohydrates, samples were heated to 100°C in 75% H_2_SO_4_ containing 2 mg/mL Anthrone reagent for 15 minutes. A set of prepared starch standards was treated in the same manner in parallel to the samples. After cooling to room temperature, sample absorbance at 578 nm was used to construct a standard curve and calculate the concentration of the samples.

For analysis of nucleic acid content, DNA was extracted from ground cells using a DNeasy Plant mini kit (Qiagen) and quantified using a Nanodrop UV spec. RNA content was assumed to be 28-fold higher than DNA content [49]. This ratio was measured in *Chlamydomonas reinhardtii*, but due to the phylogenetic similarity of the two species, and the lack of any such information for *Chromochloris*, this estimate is assumed to be sufficient.

To determine chlorophyll and total carotenoid content, ground cells were extracted with acetone twice. The pellet was saved for protein analysis, and acetone supernatant from each extraction was pooled and measured in a spectrophotometer to determine the concentrations of chlorophylls a and b, and the concentration of total carotenoids as described by Lichtenthaler and Buschmann [50].

### Determination of Model Constraints

Samples of TAP and TGP cultures were taken at various time points during growth for biomass and glucose/acetate measurements. Samples taken for glucose/acetate measurement were centrifuged at 5,000×g for three minutes, and the resulting supernatant was collected and filtered with a 0.2 μm pore size filter. For acetate analysis in TAP cultures, filtered media samples were analyzed using an acetate assay kit (Sigma MAK086) to determine extracellular acetate concentrations over time during culture growth. For glucose analysis in TGP cultures, filtered media samples were analyzed using a YSI 2900 series biochemistry analyzer. Average cell dry weight, and the change in extracellular glucose or acetate concentrations during exponential phase were used to calculate uptake rates to constrain the model.

### Metabolomics Analysis of Spent Media

Model predictions were validated with data on the extracellular lactate concentrations in TGP cultures over time during growth. This data was collected through analysis of filtered media samples on a YSI 2900 series biochemistry analyzer.

For an untargeted metabolomics analysis of spent media, samples of TGP and TAP cultures were collected in triplicate in late-exponential phase and centrifuged at 5,000×g for three minutes. The resulting supernatant was collected and filtered with a 0.2 μm pore size filter. Media samples were frozen and shipped over dry ice for analysis. Upon arrival, samples were thawed and lyophilized, then re-suspended in 500 μL of 80% methanol. Samples were then vortexed, followed by sonication for 30 min at 4 °C. Samples were incubated at -40 °C for 1 h, then vortexed and centrifuged again before 200 μL of the supernatant was collected. This sample was mixed with 5 μL of DL-o-Chlorophenylalanine (140 μg/mL) and transferred to vial for LC-MS analysis.

Sample separation was performed by a Waters 2695 LC combined with Q Exactive MS (Thermo). The LC system was comprised of ACQUITY UPLC HSS T3 (100×2.1mm×1.8 μm) with Waters 2695 LC. The mobile phase is composed of solvent A (0.05% formic acid-water) and solvent B (acetonitrile) with a gradient elution (0-1.5 min, 95-70% A; 1.5-9.5 min, 70-5% A; 9.5-13.5 min, 5% A; 13.5-13.6 min, 5-95% A; 13.6-16.0 min, 95% A). The flow rate of the mobile phase is 0.3 mL·min-1. The column temperature was maintained at 40°C, and the sample manager temperature was set at 4°C. Mass spectrometry parameters in ESI+ and ESI-mode are listed as follows: ESI+: Heater Temp 300 °C; Sheath Gas Flow rate, 45arb; Aux Gas Flow Rate, 15arb; Sweep Gas Flow Rate, 1arb; spray voltage, 3.0KV; Capillary Temp, 350 °C; S-Lens RF Level, 30%. ESI-: Heater Temp 300 °C, Sheath Gas Flow rate, 45arb; Aux Gas Flow Rate, 15arb; Sweep Gas Flow Rate, 1arb; spray voltage, 3.2KV; Capillary Temp,350 °C; S-Lens RF Level,60%. Sample peaks were identified by comparing MS spectra against the Yeast Metabolome Database (YMDB) and the Human Metabolome Database (HMDB).

### Model Robustness Analysis

An *in silico* gene essentiality analysis was performed on the genome scale metabolic model of *Chromochloris zofingiensis*. This consisted of assessing the cells growth rate for single gene deletions of every gene in the model, and assigning each gene as essential (achieve 0-50% of WT growth rate), beneficial (achieve 50-99% of WT growth rate), or non-essential (achieve 99-100% of WT growth rate). Flux Variability Analysis (FVA) was also performed on the model. Flux variability analysis (FVA)[51] is a computational technique to fully explore the solution space of a model, given a specific set of constraints. FVA produces a maximum and minimum flux for every reaction within the model, given a requirement to optimize the objective function to a certain fraction of the optimal solution. It is a key part of OptForce [52], an algorithm that evaluates which reactions must be upregulated or downregulated in order to produce a certain amount of a desired product.

## Notes

### Competing Interest Statement

The authors have declared no competing interest.

### Summary of Updates

we quantified the production of lactate in the cell and revised the flux maps to match

