## Supplemental Material for "Genome-Scale Metabolic Model Accurately Predicts Fermentation of Glucose by *Chromochloris zofingiensis*"

**Supplemental Information**

**Supplemental File 1:iCre1915** – Genome scale metabolic model of *Chromochloris zofingiensis* in SBML format.

**Supplemental File 2: Metabolites** –List of all metabolites included in iCre1915.

**Supplemental File 3: Reactions**- List of all reactions included in iCre1915, including genes associated with each reaction.

**Supplemental File 4: MEMOTE Results-** MEMOTE analysis summary.

**Supplemental File 5: Metabolomics data**- All metabolomics data from the analysis of spent media in TGP and TAP cultures, used for confirmation of the formation of fermentation products.

**Supplemental File 6: FVA results, Biomass case-** Full results of flux variability analysis in production of biomass.

**Supplemental File 7: FVA results, Astaxanthin full biomass case-** Full results of flux variability analysis, examining astaxanthin production potential with production of biomass.

**Supplemental File 8: FVA results, Astaxanthin no biomass case-** Full results of flux variability analysis, examining astaxanthin production potential with no production of biomass.

**Supplemental File 9: FVA results, Lipids full biomass case-** Full results of flux variability analysis, examining lipid production potential with production of biomass.

**Supplemental File 10: FVA results, Lipids no biomass case-** Full results of flux variability analysis, examining lipid production potential with no production of biomass.

**Supplemental Tables and Figures:**

Table S1. List of reactions manually added to RAPS output to construct genome scale metabolic model of *Chromochloris zofingiensis.*

Table S2. List of reactions manually removed from RAPS output to generate genome-scale metabolic model of *Chromochloris zofingiensis.*

Figure S1. Growth data for C. zofingiensis in each media type.

Figure S2. Acetate and glucose consumption in TAP and TGP cultures.

Figure S3. YSI data on extracellular metabolites in TGP cultures.

Table S3. Constraints imposed for flux balance analysis of *C. zofingiensis* genome-scale model.

Table S4. Biomass formation equation for each growth condition.

Table S5 List of essential genes for each growth condition

**<u>Table S1.</u> List of reactions manually added to RAPS output to construct genome scale metabolic model of *Chromochloris zofingiensis*.** KEGG enzyme identification numbers are listed where applicable.

| Reaction ID | EC Number | Enzyme Name | Reaction |
|---|---|---|---|
| NanoG1428 | 1.17.7.1 | (E)-4-hydroxy-3-methylbut-2-enyl-diphosphate synthase | 2 Reduced ferredoxin + H + 2-C-Methyl-D-erythritol 2,4-cyclodiphosphate --> 2 Oxidized ferredoxin + 1-Hydroxy-2-methyl-2-butenyl 4-diphosphate + H2O |
| DMORm | 1.1.1.86 | 2,3-Dihydroxy-3-methylbutanoate:NADP+ oxidoreductase (isomerizing), mitochondria | NADPH + H + (S)-2-Acetolactate --> (R)-2,3-Dihydroxy-3-methylbutanoate + NADP |
| NanoG1326 | 2.5.1.78 | 6,7-dimethyl-8-ribityllumazine synthase | L-3,4-Dihydroxybutan-2-one 4-phosphate + 5-Amino-6-(D-ribitylamino)uracil --> H+ Phosphate + 6,7-Dimethyl-8-(1-D-ribityl)lumazine + 2 H2O |
| ATPSh | 3.6.3.14 | ATP synthase | 4 H + Phosphate + ADP --> 3 H + H2O + ATP |
| ATPSm | 3.6.3.14 | F0F1-ATP synthase Complex V | 3 H+ Phosphate + ADP --> 2 H + H2O + ATP |
| NADHOR | 1.6.5.3 | NADH:ubiquinone oxidoreductase Complex I | NADH + 5 H + Ubiquinone --> NAD + 4 H + Ubiquinol |
| PNORm | 1.1.1.169 | Pantoate:NADP+ 2-oxidoreductase, mitochondria | (R)-Pantoate + NADP <=> NADPH + H + 2-Dehydropantoate |
| PBALm | 6.3.2.1 | Pantoate:beta-alanine ligase (AMP-forming) | (R)-Pantoate + beta-Alanine + ATP --> (R)-Pantothenate + 3 H + AMP + Diphosphate |
| NanoG0910 | 3.1.3.4 | Phosphatidate phosphatase (n-C20 5) | Phosphatidate (20:5) + H2O --> 1,2-Diacyl-sn-glycerol (C20:5) + Phosphate |
| TRPth | | Tryptophan permease, cytosol to chloroplast transport reaction | H + L-Tryptophan_c <=> H + L-Tryptophan_h |
| RETACIH | 3.1.1.64 | all-trans-retinyl ester isomerohydrolase (acetate-utilizing) | all-trans-Retinyl acetate + H2O --> 11-cis-Retinol + H + Acetate |

| | | | |
|---|---|---|---|
| NanoG1070 | 3.6.1.26 | cdp-diacylglycerol phosphatidylhydrolase | H2O + CDP-diacylglycerol (n-c20:5) --> H + CMP + Phosphatidate (20:5) |
| NanoG1648 | 2.7.8.11 | cdp-diacylglycerol---inositol 3-phosphatidyltransferase | myo-Inositol + CDP-diacylglycerol (n-c20:4) --> phosphatidylinositol (C20:4) + H + CMP |
| CBFC | 1.10.99.1 | cytochrome b6/f complex | 2 Plastocyanin(Cu2+) + 2 H + reduced plastoquinone --> 2 Plastocyanin(Cu+) + 4 H + oxidized plastoquinone |
| CYOO6m | 1.9.3.1 | cytochrome c oxidase Complex IV | 4 Ferrocytochrome + 8 H + O2 --> 4 Ferricytochrome + 4 H + 2 H2O |
| NanoG0676 | 1.14.19.6 | delta 6 desaturase | 2 Ferrocytochrome b5 + 2 H + O2 + (9Z,12Z,15Z)-Octadecatrienoyl-CoA --> 2 Ferricytochrome b5 + (6Z,9Z,12Z,15Z)-octadecatetraenoyl-CoA + 2 H2O |
| NanoG0864 | 2.3.1.199 | fatty acid elongase | (6Z,9Z,12Z,15Z)-octadecatetraenoyl-CoA + H + Malonyl-CoA --> CO2 + Coenzyme A + C20:(4) 3- oxoacylCoA |
| NanoG1190 | 3.1.2.2 | fatty acid elongase | H + Malonyl-CoA + gamma-linolenoyl-CoA --> CO2 + 3-oxo-eicosatrienoyl-CoA + Coenzyme A |
| NanoG0999 | 2.3.1.15 | glycerol-3-phosphate: acyl-coa acyltransferase C20:4 | Glycerol 3-phosphate + (8Z,11Z,14Z,17Z)-Icosatetraenoyl-CoA (n-c20:4) --> 1-(9Z,12Z,15Z,18Z)-Eicosapentaenoyl-sn-glycerol-3-phosphate + Coenzyme A |
| NanoG1001 | 2.3.1.51 | glycerol-3-phosphate: acyl-coa acyltransferase C20:5 | Glycerol 3-phosphate + (5Z,8Z,11Z,14Z,17Z)-Icosapentaenoyl-CoA --> 1-(5Z,8Z,11Z,14Z,17Z)-Eicosapentaenoyl-sn-glycerol-3-phosphate + Coenzyme a |
| IMGPS | 4.3.2.10 | imizadole glycerol phosphate synthase, cytosol | 5-_(5-phospho-1-deoxyribulos-1-ylamino)methylideneamino_-1-(5-phosphoribosyl)imidazole-4-carboxamide + L-Glutamine --> H + D-erythro-1-(Imidazol-4-yl)glycerol 3-phosphate + 5-Amino-1-(5-Phospho-D-ribosyl)imidazole-4-carboxamide + L-Glutamate |
| NanoG0263 | 1.1.1.86 | ketol-acid reductoisomerase | NADPH + H + (S)-2-Aceto-2-hydroxybutanoate_h --> (R)-2,3-Dihydroxy-3-methylpentanoate + NADP |
| PSIblue | | photosystem I (blue light-activated) | Oxidized ferredoxin + 2 Plastocyanin(Cu+) + H + 2 photon (406 to 454 nm, indigo/blue) --> Reduced ferredoxin + 2 Plastocyanin(Cu2+) |
| PSIred | | photosystem I (red light-activated) | Oxidized ferredoxin + 2 Plastocyanin(Cu+) + 2 H + 2 photon (662 to 691 nm, red) --> Reduced ferredoxin + 2 Plastocyanin(Cu2+) |

| | | | |
|---|---|---|---|
| PSIIblue | | photosystem II (blue light-activated) | 2 Oxidized plastoquinone + 2 H2O + 4 photon (378 to 482 nm, violet/blue) --> O2 + 2 reduced plastoquinone |
| PSIIred | | photosystem II (red light-activated) | 2 oxidized plastoquinone + 2 H2O + 4 photon (659 to 684 nm, red) --> O2 + 2 reduced plastoquinone |
| RBPCh | 4.1.1.39 | ribulose-bisphosphate carboxylase | CO2 + H2O + D-Ribulose 1,5-bisphosphate --> 2 H + 2 3-Phospho-D-glycerate |
| CYOR_q8_m | 1.10.2.2 | ubiquinol-cytochrome c oxidoreductase Complex III | 2 Ferricytochrome c + 2 H + Ubiquinol --> 2 Ferrocytochrome c + 4 H + Ubiquinone |
| ZAC | 1.14.99.64 | Adonixanthin production | Zeaxanthin + H + NADPH + O2 --> Adonixanthin + H2O + NADP |
| AAxC | 1.14.99.64 | Astaxanthin production | Adonixanthin + H + NADPH + O2 --> Astaxanthin + H2O + NADP |

**Table S2. List of reactions manually removed from RAPS output to generate genome-scale metabolic model of *Chromochloris zofingiensis.*** KEGG enzyme identification numbers are listed where applicable.

| Reaction ID | EC Number | Enzyme | Reaction |
|---|---|---|---|
| NanoG1423 | 2.7.7.60 | 2-c-methyl-d-erythritol 4-phosphate cytidylyltransferase | H + CTP + 2-C-Methyl-D-erythritol 4-phosphate --> Diphosphate + 4-(Cytidine 5'-diphospho)-2-C-methyl-D-erythritol |
| NanoG0467 | 2.3.1.61 | 2-oxoglutarate dehydrogenase E2 component | S-Succinyldihydrolipoamide + Coenzyme A --> Dihydrolipoamide + Succinyl-CoA |
| NanoG0615 | 3.6.3.14 | ATP synthase | 4 H + Phosphate + ADP --> 3 H + H2O + ATP |
| HYPS_13bdg_LPAREN_c_RPAREN | | Chrysolaminarin Hypothetical storage | 1,3-beta-D-Glucan (Callose) --> |
| Tr_DGTPt_m | | Deoxynucleotide carrier (dgtp), mitochondrial | dGDP + dGTP --> 2-Deoxyguanosine 5-diphosphate + dGTP |
| NanoG0606 | | F0F1-ATP synthase | H + Phosphate + ADP --> 2 H + H2O + ATP |

| | | | |
|---|---|---|---|
| | | Complex V | |
| M5TRPI | 5.3.1.23 | S-methyl-5-thioribose-1-phosphate isomerase | --> S-Methyl-5-deoxy-D-ribose 1-phosphate |
| NanoG1493 | 3.5.3.4 | allantoicase | Allantoate + H2O--> (-)-Ureidoglycolate + Urea |
| NanoG0608 | 7.1.1.9 | cytochrome-c oxidase | 4 Ferrocytochrome c + 4 H + O2 --> 4 Ferricytochrome c + 2 H2O |
| DM_o2D_u | | demand removing dummy o2 from system | O2 --> sink |
| NanoG1539 | 3.5.2.3 | dihydroorotase | H + N-Carbamoyl-L-aspartate --> (S)-Dihydroorotate + H2O |
| DVPCHLD450OR | 1.3.1.33 | divinylprotochlorophyllide oxidoreductase (light-dependent, 450 nm) | NADPH + H + Divinylprotochlorophyllide a + photon (417 to 472 nm, indigo/blue) --> Divinyl chlorophyllide a + NADP |
| NanoG1585 | 1.12.7.2 | ferredoxin hydrogenase | Reduced ferredoxin + 2 H --> Oxidized ferredoxin + H2 |
| EX_glc_DASH_d_LPARE N_e_RPAREN | | glucose exchange | beta-D-Glucose --> sink |
| NanoG1239 | 1.5.1.5 | methylenetetrahydrofolate dehydrogenase (nadp+) | 5,10-Methylenetetrahydrofolate + NADP --> NADPH + 5,10-Methenyltetrahydrofolate |
| NanoG0405, NanoG0406, NanoG0407 | 4.2.1.11 | phosphopyruvate hydratase | H + D-Glycerate 2-phosphate --> Phosphoenolpyruvate + H2O |
| NanoG0607 | 7.1.1.8 | quinol-cytochrome-c reductase | Ferricytochrome c + ubiquinol --> 2 Ferrocytochrome c + 2 H + ubiquinone |
| NanoG0093 | | s-methyl-5-thioribose-1-phosphate isomerase | --> S-Methyl-5-thio-alpha-D-ribose 1-phosphate |
| SUCL_gdp_m | 6.2.1.4 | succinyl-CoA ligase (GDP-forming) | Succinate + Coenzyme A + GTP --> Phosphate + Guanosine 5-diphosphate + Succinyl-CoA |

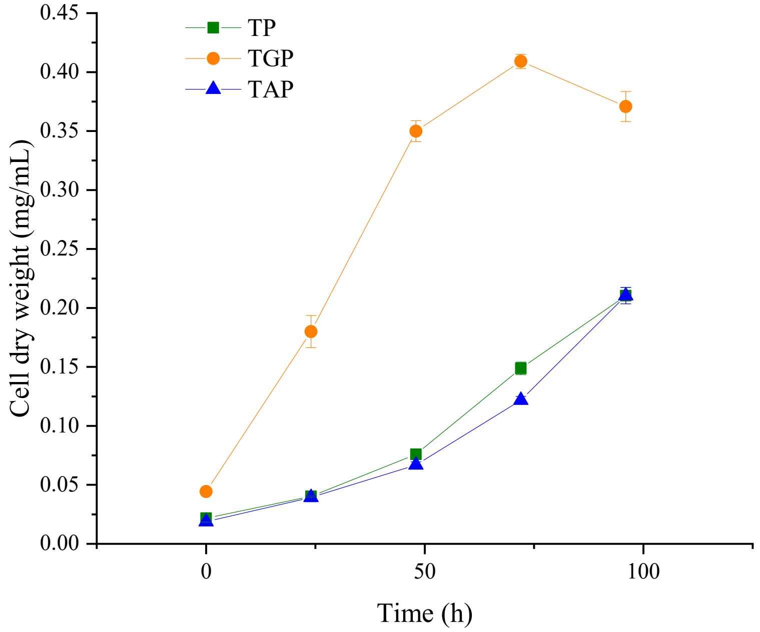


**<u>Figure S1</u>. Growth data for C. zofingiensis in each media type.** Data presented here was used to calculate growth rates used during pFBA simulations for autotrophic growth (TP), mixotrophic growth on acetate (TAP), and heterotrophic

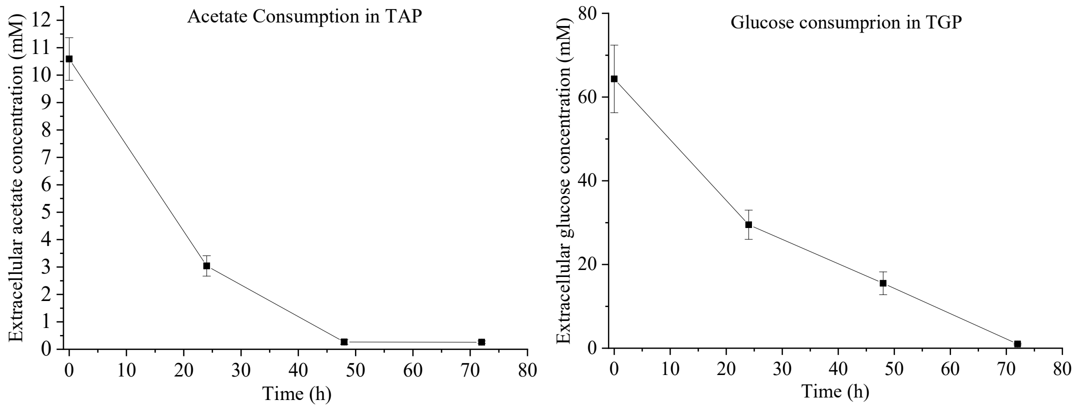


**<u>Figure S2.</u> Acetate and glucose consumption in TAP and TGP cultures**.
Data shown here was used to calculate acetate and glucose uptake flux values

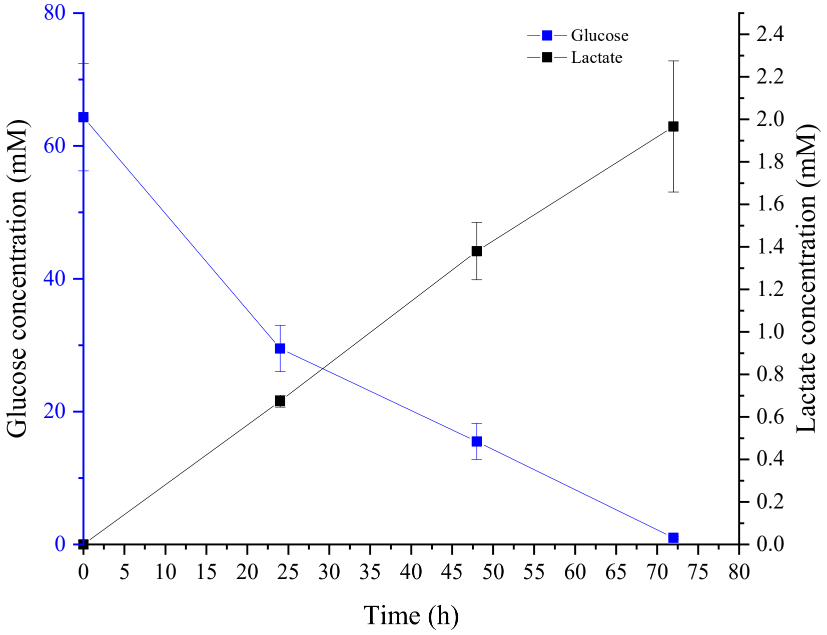


**Figure S3. YSI data on extracellular metabolites in TGP cultures.** Glucose consumption and lactate excretion were measured.

**Table S3. Constraints imposed for flux balance analysis of *C. zofingiensis* genome-scale model.**

| Model Constraints (mmol/gDW-hr) | TP | TAP | TGP |
|---|---|---|---|
| Water, maximum | 1000 | 1000 | 1000 |
| Ammonia, maximum | 1000 | 1000 | 1000 |
| Sulfate, maximum | 1000 | 1000 | 1000 |
| Phosphate, maximum | 1000 | 1000 | 1000 |
| Oxygen, maximum | 1000 | 1000 | 1000 |
| Magnesium, maximum | 1 | 1 | 1 |
| Iron, maximum | 1 | 1 | 1 |
| Cobalt, maximum | 1 | 1 | 1 |

| | | | |
|---|---|---|---|
| Growth Rate, fixed | 0.0264 | 0.0274 | 0.042 |
| Carbon Dioxide, maximum | 0.9728 | 0.9728 | 0.9728 |
| Photons, maximum | 43.98 | 43.98 | 43.98 |
| Acetate, fixed | 0 | 3.22 | 0 |
| Glucose, fixed | 0 | 0 | 2.41 |

**Table S4. Biomass formation equation for each growth condition**. Tris-phosphate medium (TP) is used for photoautotrophic growth, trisacetate phosphate medium (TAP) is used for mixotrophic growth and tris-glucose phosphate (TGP) medium is used for heterotrophic growth.

| Category | Model ID | TP | TAP | TGP |
|---|---|---|---|---|
| energy | "atp_c" | 92.40009 | 92.4153 | 92.40436 |
| misc | "h2o_c" | 92.4 | 92.4 | 92.4 |
| protein | "alatrna_c" | 0.147471 | 0.059995 | 0.046893 |
| protein | "argtrna_c" | 0.113543 | 0.148301 | 0.064127 |
| protein | "asntrna_c" | 0.285119 | 0.282816 | 0.07935 |
| protein | "asptrna_c" | 0.285119 | 0.282816 | 0.07935 |
| protein | "cystrna_c" | 0.103309 | 0.119425 | 0.017733 |
| protein | "glntrna_c" | 0.274907 | 0.367701 | 0.07965 |
| protein | "glutrna_c" | 0.274907 | 0.367701 | 0.07965 |
| protein | "glytrna_c" | 0.159853 | 0.020424 | 0.040534 |
| protein | "histrna_c" | 0.199687 | 0.230838 | 0.025467 |
| protein | "iletrna_c" | 0.070988 | 0.142402 | 0.033298 |
| protein | "leutrna_c" | 0.330941 | 0.668692 | 0.135396 |
| protein | "lystrna_c" | 0.120682 | 0.046467 | 0.065531 |
| protein | "mettrna_c" | 0.005163 | 0.006816 | 0.002887 |
| protein | "phetrna_c" | 0.09776 | 0.133263 | 0.031589 |
| protein | "protrna_c" | 0.068857 | 0.0847 | 0.02495 |
| protein | "sertrna_c" | 0.212073 | 0.139757 | 0.071145 |
| protein | "thrtrna_c" | 0.051672 | 0.059733 | 0.00887 |
| protein | "trptrna_c" | 0.101136 | 0.116913 | 0.012898 |
| protein | "tyrtrna_c" | 0.047671 | 0.052072 | 0.010803 |
| protein | "valtrna_c" | 0.118139 | 0.066489 | 0.054286 |
| nucleic acid | "datp_c" | 0.000834 | 0.00056 | 0.00016 |
| nucleic acid | "dctp_c" | 0.000868 | 0.000583 | 0.000166 |
| nucleic acid | "dgtp_c" | 0.000868 | 0.000583 | 0.000166 |
| nucleic acid | "dttp_c" | 0.000834 | 0.00056 | 0.00016 |
| energy | "ctp_c" | 9.47E-05 | 0.015921 | 0.004537 |
| energy | "gtp_c" | 9.47E-05 | 0.015921 | 0.004537 |
| energy | "utp_c" | 9.10E-05 | 0.015297 | 0.004359 |
| carbohydrates | "starch300_h" | 0.000447 | 0.000441 | 0.003678 |
| carbohydrates | "man_c" | 0.164305 | 0.108079 | 0.349895 |
| carbohydrates | "arab_L_c" | 0.262268 | 0.172518 | 0.558512 |
| carbohydrates | "gal_c" | 0.34876 | 0.229412 | 0.742702 |
| lipids | "mgdg1839Z12Z15Z1644Z7Z10Z13Z_h" | 0.004361 | 0.000199 | 0.000144 |
| lipids | "mgdg1839Z12Z15Z1637Z10Z13Z_h" | 0.01286 | 0.00205 | 0.000729 |

| | | | | |
|---|---|---|---|---|
| lipids | "mgdg1839Z12Z15Z1634Z7Z10Z_h" | 0.01286 | 0.00205 | 0.000729 |
| lipids | "mgdg1829Z12Z1644Z7Z10Z13Z_h" | 0.02296 | 0.018435 | 0.060642 |
| lipids | "mgdg1829Z12Z1637Z10Z13Z_h" | 0.001543 | 0.006133 | 0.004162 |
| lipids | "mgdg1829Z12Z1634Z7Z10Z_h" | 0.001543 | 0.006133 | 0.004162 |
| lipids | "mgdg1839Z12Z15Z1627Z10Z_h" | 0.011255 | 0.000228 | 0.000136 |
| lipids | "mgdg1829Z12Z1627Z10Z_h" | 0.010919 | 0.032837 | 0.002786 |
| lipids | "mgdg1829Z12Z1617Z_h" | 0.000642 | 0.002465 | 0.001792 |
| lipids | "mgdg1829Z12Z1619Z_h" | 0.000642 | 0.002464 | 0.001792 |
| lipids | "mgdg1839Z12Z15Z160_h" | 0.000616 | 0.00054 | 0.00025 |
| lipids | "mgdg1829Z12Z160_h" | 0.000686 | 0.006722 | 0.001026 |
| lipids | "dgdg1839Z12Z15Z1644Z7Z10Z13Z_h" | 0.000826 | 0.00317 | 0.000894 |
| lipids | "dgdg1839Z12Z15Z1637Z10Z13Z_h" | 0.002492 | 0.015918 | 0.00267 |
| lipids | "dgdg1839Z12Z15Z1634Z7Z10Z_h" | 0.002492 | 0.009108 | 0.002487 |
| lipids | "dgdg1839Z12Z15Z1627Z10Z_h" | 0.014009 | 0.000228 | 0.000136 |
| lipids | "dgdg1829Z12Z1637Z10Z13Z_h" | 0.000911 | 0.003555 | 0.002514 |
| lipids | "dgdg1829Z12Z1634Z7Z10Z_h" | 0.000911 | 0.003554 | 0.002514 |
| lipids | "dgdg1829Z12Z1627Z10Z_h" | 0.004792 | 0.00258 | 0.000658 |
| lipids | "dgdg1819Z1637Z10Z13Z_h" | 0.000498 | 0.000401 | 0.002227 |
| lipids | "dgdg1819Z1634Z7Z10Z_h" | 0.000498 | 0.000401 | 0.002228 |
| lipids | "dgdg1819Z1627Z10Z_h" | 0.006706 | 0.000903 | 0.007644 |
| lipids | "dgdg1839Z12Z15Z160_h" | 0.015413 | 0.000222 | 0.000138 |
| lipids | "dgdg1829Z12Z1617Z_h" | 0.001049 | 0.004125 | 0.002881 |
| lipids | "dgdg1829Z12Z1619Z_h" | 0.001049 | 0.004197 | 0.002881 |
| lipids | "dgdg1829Z12Z160_h" | 0.04152 | 0.001196 | 0.000526 |
| lipids | "dgdg1819Z1617Z_h" | 0.000667 | 1.44E-06 | 0.003589 |
| lipids | "dgdg1819Z1619Z_h" | 0.000667 | 1.45E-06 | 0.003588 |
| lipids | "dgdg1819Z160_h" | 0.001495 | 0.003391 | 0.004971 |
| lipids | "dgts1839Z12Z15Z1845Z9Z12Z15Z_c" | 0.00128 | 0.000381 | 0.000245 |
| lipids | "dgts1839Z12Z15Z1835Z9Z12Z_c" | 0.003841 | 0.000232 | 0.000143 |
| lipids | "dgts1829Z12Z1845Z9Z12Z15Z_c" | 0.000226 | 0.000843 | 0.000646 |
| lipids | "dgts1829Z12Z1835Z9Z12Z_c" | 0.010164 | 0.008348 | 0.001199 |
| lipids | "dgts1819Z1845Z9Z12Z15Z_c" | 0.00062 | 0.000128 | 0.003164 |
| lipids | "dgts18111Z1845Z9Z12Z15Z_c" | 0.00062 | 0.00013 | 0.003162 |
| lipids | "dgts1601845Z9Z12Z15Z_c" | 0.012971 | 0.000769 | 0.000411 |
| lipids | "dgts1829Z12Z1829Z12Z_c" | 0.002015 | 0.006851 | 0.004592 |
| lipids | "dgts1839Z12Z15Z1819Z_c" | 0.000722 | 0.001723 | 0.001829 |
| lipids | "dgts1829Z12Z1819Z_c" | 0.000216 | 0.000463 | 0.000723 |
| lipids | "dgts1839Z12Z15Z18111Z_c" | 0.000722 | 0.001723 | 0.001829 |
| lipids | "dgts1829Z12Z18111Z_c" | 0.000216 | 0.000462 | 0.000723 |
| lipids | "dgts1819Z1835Z9Z12Z_c" | 0.000808 | 0.00181 | 0.002 |
| lipids | "dgts18111Z1835Z9Z12Z_c" | 0.000808 | 0.001827 | 0.002 |

| | | | | |
|---|---|---|---|---|
| lipids | "dgts1601835Z9Z12Z_c" | 0.050541 | 0.000223 | 0.000138 |
| lipids | "dgts1819Z1829Z12Z_c" | 0.000584 | 0.000216 | 0.002452 |
| lipids | "dgts18111Z1829Z12Z_c" | 0.000584 | 0.000215 | 0.002452 |
| lipids | "dgts1819Z1819Z_c" | 0.000193 | 0.000252 | 0.000765 |
| lipids | "dgts1819Z18111Z_c" | 0.000193 | 0.000252 | 0.000765 |
| lipids | "dgts18111Z1819Z_c" | 0.000193 | 0.000252 | 0.000765 |
| lipids | "dgts18111Z18111Z_c" | 0.000193 | 0.000255 | 0.000765 |
| lipids | "dgts1601829Z12Z_c" | 0.009194 | 0.001076 | 0.000478 |
| lipids | "dgts1601819Z_c" | 0.00022 | 0.000117 | 0.000656 |
| lipids | "dgts16018111Z_c" | 0.00022 | 0.000119 | 0.000656 |
| lipids | "sqdg1839Z12Z15Z160_h" | 0.002929 | 0.000226 | 0.000138 |
| lipids | "sqdg1829Z12Z160_h" | 0.004429 | 0.001016 | 0.00047 |
| lipids | "sqdg1819Z160_h" | 0.001022 | 0.002953 | 0.003177 |
| lipids | "sqdg18111Z160_h" | 0.001022 | 0.003028 | 0.003175 |
| lipids | "sqdg160_h" | 0.016608 | 0.077682 | 0.000444 |
| lipids | "asqdpa1819Z160_c" | 0.001494 | 0.000985 | 0.004733 |
| lipids | "asqdpa18111Z160_c" | 0.001494 | 0.000985 | 0.004732 |
| lipids | "asqdpa1829Z12Z160_c" | 0.010284 | 0.001092 | 0.00048 |
| lipids | "asqdpa1839Z12Z15Z160_c" | 0.005282 | 0.000212 | 0.000138 |
| lipids | "asqdca1819Z160_c" | 0.001495 | 0.003243 | 0.004754 |
| lipids | "asqdca18111Z160_c" | 0.001495 | 0.003397 | 0.004738 |
| lipids | "asqdca1829Z12Z160_c" | 0.010318 | 0.001092 | 0.00048 |
| lipids | "asqdca1839Z12Z15Z160_c" | 0.005296 | 0.000247 | 0.000138 |
| lipids | "pg1839Z12Z15Z1613E_h" | 0.004541 | 0.088477 | 0.002187 |
| lipids | "pg1839Z12Z15Z160_h" | 0.002733 | 0.000206 | 0.000138 |
| lipids | "pg1829Z12Z1613E_h" | 0.024139 | 0.020967 | 0.092887 |
| lipids | "pg1829Z12Z160_h" | 0.004518 | 0.001009 | 0.00047 |
| lipids | "pg1819Z1613E_h" | 0.000856 | 0.001659 | 0.005857 |
| lipids | "pg18111Z1613E_h" | 0.000856 | 0.001659 | 0.005857 |
| lipids | "pg1819Z160_h" | 0.001171 | 0.003122 | 0.003644 |
| lipids | "pg18111Z160_h" | 0.001171 | 0.003234 | 0.003644 |
| lipids | "pe1829Z12Z1835Z9Z12Z_c" | 0.000629 | 0.003535 | 0.00089 |
| lipids | "pe1819Z1845Z9Z12Z15Z_c" | 0.000189 | 1.11E-05 | 4.82E-05 |
| lipids | "pe1819Z1835Z9Z12Z_c" | 0.000265 | 0.000843 | 0.000751 |
| lipids | "pe18111Z1845Z9Z12Z15Z_c" | 0.000344 | 0.000508 | 0.001319 |
| lipids | "pe18111Z1835Z9Z12Z_c" | 0.000525 | 0.001896 | 0.002162 |
| lipids | "pe1801845Z9Z12Z15Z_c" | 0.002127 | 0.000172 | 0.000144 |
| lipids | "pe1801835Z9Z12Z_c" | 0.009102 | 0.000188 | 0.000144 |
| lipids | "pail18111Z160_c" | 0.001451 | 0.003345 | 0.004934 |
| lipids | "pail1819Z160_c" | 0.000652 | 0.002248 | 0.002016 |
| lipids | "tag16018111Z160_c" | 0.001446 | 0.003191 | 0.000982 |

| | | | | |
|---|---|---|---|---|
| lipids | "tag1601819Z160_c" | 0.001444 | 0.003179 | 0.000987 |
| lipids | "tag1801819Z160_c" | 0.001511 | 0.001004 | 0.000848 |
| lipids | "tag18111Z18111Z160_c" | 0.000709 | 0.000572 | 0.005096 |
| lipids | "tag18111Z1819Z160_c" | 0.000709 | 0.000572 | 0.005096 |
| lipids | "tag1819Z18111Z160_c" | 0.000709 | 0.000572 | 0.005096 |
| lipids | "tag1819Z1819Z160_c" | 0.00071 | 0.000572 | 0.005102 |
| lipids | "tag16018111Z180_c" | 0.001511 | 0.001004 | 0.000849 |
| lipids | "tag1601819Z180_c" | 0.001511 | 0.001 | 0.000847 |
| lipids | "tag1801819Z180_c" | 0.001219 | 0.000454 | 0.000759 |
| lipids | "tag18111Z18111Z180_c" | 0.000929 | 0.000157 | 0.019937 |
| lipids | "tag18111Z1819Z180_c" | 0.000929 | 0.000157 | 0.019946 |
| lipids | "tag1819Z18111Z180_c" | 0.000929 | 0.000157 | 0.019923 |
| lipids | "tag1819Z1819Z180_c" | 0.000151 | 0.000195 | 0.001147 |
| lipids | "tag16018111Z18111Z_c" | 0.000709 | 0.000572 | 0.005097 |
| lipids | "tag1601819Z18111Z_c" | 0.000709 | 0.000572 | 0.005097 |
| lipids | "tag1801819Z18111Z_c" | 0.000319 | 0.000157 | 0.002389 |
| lipids | "tag18111Z18111Z18111Z_c" | 0.000126 | 0.000228 | 0.001182 |
| lipids | "tag18111Z1819Z18111Z_c" | 0.000123 | 0.000234 | 0.001159 |
| lipids | "tag1819Z18111Z18111Z_c" | 0.000123 | 0.000234 | 0.001158 |
| lipids | "tag1819Z1819Z18111Z_c" | 0.000119 | 0.000422 | 0.001132 |
| lipids | "tag16018111Z1819Z_c" | 0.000709 | 0.000572 | 0.005097 |
| lipids | "tag1601819Z1819Z_c" | 0.000712 | 0.000572 | 0.005122 |
| lipids | "tag1801819Z1819Z_c" | 0.000319 | 0.000157 | 0.002388 |
| lipids | "tag18111Z18111Z1819Z_c" | 0.000123 | 0.000234 | 0.001159 |
| lipids | "tag18111Z1819Z1819Z_c" | 0.000123 | 0.000234 | 0.001159 |
| lipids | "tag1819Z18111Z1819Z_c" | 0.000123 | 0.000234 | 0.001159 |
| lipids | "tag1819Z1819Z1819Z_c" | 0.000119 | 0.000239 | 0.001131 |
| lipids | "tag16018111Z1835Z9Z12Z_c" | 0.001467 | 0.001218 | 0.000561 |
| lipids | "tag1601819Z1835Z9Z12Z_c" | 0.001466 | 0.001213 | 0.000563 |
| lipids | "tag1801819Z1835Z9Z12Z_c" | 0.001232 | 0.000868 | 0.000781 |
| lipids | "tag18111Z18111Z1835Z9Z12Z_c" | 0.000237 | 0.000563 | 0.001318 |
| lipids | "tag18111Z1819Z1835Z9Z12Z_c" | 0.000179 | 0.000531 | 0.001079 |
| lipids | "tag1819Z18111Z1835Z9Z12Z_c" | 0.000175 | 0.000529 | 0.001066 |
| lipids | "tag1819Z1819Z1835Z9Z12Z_c" | 0.000761 | 0.000495 | 0.003517 |
| lipids | "tag16018111Z1845Z9Z12Z15Z_c" | 0.001512 | 0.003234 | 0.004827 |
| lipids | "tag1601819Z1845Z9Z12Z15Z_c" | 0.001512 | 0.003371 | 0.004831 |
| lipids | "tag1801819Z1845Z9Z12Z15Z_c" | 0.001229 | 0.000351 | 0.007405 |
| lipids | "tag18111Z18111Z1845Z9Z12Z15Z_c" | 0.000149 | 0.000174 | 0.001226 |
| lipids | "tag18111Z1819Z1845Z9Z12Z15Z_c" | 0.000149 | 0.000174 | 0.001225 |
| lipids | "tag1819Z18111Z1845Z9Z12Z15Z_c" | 0.000149 | 0.000174 | 0.001225 |
| lipids | "tag1819Z1819Z1845Z9Z12Z15Z_c" | 0.000149 | 0.000174 | 0.001228 |

| | | | | |
|---|---|---|---|---|
| misc | "ac_c" | 0.03709 | 0.024398 | 0.078985 |
| misc | "ppa_c" | 0.030069 | 0.019779 | 0.064033 |
| misc | "but_c" | 0.025282 | 0.016631 | 0.05384 |
| misc | "glyc_c" | 0.012093 | 0.007954 | 0.025752 |
| pigments | "chla_u" | 0.029764 | 0.030463 | 0.000732 |
| pigments | "chlb_u" | 0.013049 | 0.010297 | 0.000306 |
| pigments | "rhodopsin_s" | 1.60E-07 | 1.04E-06 | 1.04E-06 |
| pigments | "acaro_h" | 7.76E-05 | 0.000504 | 0.000504 |
| pigments | "anxan_u" | 1.55E-05 | 0.000101 | 0.000101 |
| pigments | "caro_u" | 0.000217 | 0.001412 | 0.001412 |
| pigments | "loroxan_u" | 0.000101 | 0.000655 | 0.000655 |
| pigments | "lut_u" | 0.000194 | 0.00126 | 0.00126 |
| pigments | "neoxan_u" | 8.53E-05 | 0.000555 | 0.000555 |
| pigments | "vioxan_u" | 5.43E-05 | 0.000353 | 0.000353 |
| pigments | "zaxan_u" | 4.65E-05 | 0.000303 | 0.000303 |
| energy | "nad_c" | 0.041244 | 0.001787 | 0.001787 |
| energy | "nadh_c" | 0.001039 | 0.000045 | 0.000045 |
| energy | "nadp_c" | 0.002585 | 0.000112 | 0.000112 |
| energy | "nadph_c" | 0.007732 | 0.000335 | 0.000335 |
| misc | "btn_c" | 2.22E-05 | 2.56E-05 | 7.08E-06 |
| misc | "thmmp_c" | 0.00248 | 0.002851 | 0.000789 |
| energy | "fad_c" | 0.00248 | 0.002851 | 0.000789 |
| misc | "gthrd_c" | 0.000656 | 0.000754 | 0.000209 |
| | "adp_c" | -92.4 | -92.4 | -92.4 |
| | "h_c" | -92.4 | -92.4 | -92.4 |
| | "pi_c" | -92.4 | -92.4 | -92.4 |
| | "trnaala_c" | -0.14747 | -0.06 | -0.04689 |
| | "trnaarg_c" | -0.11354 | -0.1483 | -0.06413 |
| | "trnaasn_c" | -0.28512 | -0.28282 | -0.07935 |
| | "trnaasp_c" | -0.28512 | -0.28282 | -0.07935 |
| | "trnacys_c" | -0.10331 | -0.11942 | -0.01773 |
| | "trnagln_c" | -0.27491 | -0.3677 | -0.07965 |
| | "trnaglu_c" | -0.27491 | -0.3677 | -0.07965 |
| | "trnagly_c" | -0.15985 | -0.02042 | -0.04053 |
| | "trnahis_c" | -0.19969 | -0.23084 | -0.02547 |
| | "trnaile_c" | -0.07099 | -0.1424 | -0.0333 |
| | "trnaleu_c" | -0.33094 | -0.66869 | -0.1354 |
| | "trnalys_c" | -0.12068 | -0.04647 | -0.06553 |
| | "trnamet_c" | -0.00516 | -0.00682 | -0.00289 |
| | "trnaphe_c" | -0.09776 | -0.13326 | -0.03159 |
| | "trnapro_c" | -0.06886 | -0.0847 | -0.02495 |

| | | | | |
|---|---|---|---|---|
| | "trnaser_c" | -0.21207 | -0.13976 | -0.07114 |
| | "trnathr_c" | -0.05167 | -0.05973 | -0.00887 |
| | "trnatrp_c" | -0.10114 | -0.11691 | -0.0129 |
| | "trnatyr_c" | -0.04767 | -0.05207 | -0.0108 |
| | "trnaval_c" | -0.11814 | -0.06649 | -0.05429 |

**<u>Table S5</u> List of essential genes for each growth condition** (E = essential, NE = non-essential).

| Gene | Name | TP | TAP | TGP | Notes/Associated Reactions |
|---|---|---|---|---|---|
| Cz04g39260.t1 | | E | E | E | glutamate-5-semialdehyde dehydrogenase |
| Cz15g07080.t1 | | E | E | E | argininosuccinate lyase |
| Cz18g04260.t1 | | E | E | E | argininosuccinate synthase |
| Cz12g10090.t1 | | E | E | E | 2-C-methyl-D-erythritol 4-phosphate cytidylyltransferase |
| Cz07g19130.t1 | | E | E | E | 1-deoxy-D-xylulose-5-phosphate reductoisomerase |
| Cz02g35110.t1 | | E | E | E | Geranylgeranyl reductases |
| Cz11g24270.t1 | | E | E | E | 4-(cytidine 5'-diphospho)-2-C-methyl-D-erythritol kinase |
| Cz02g35280.t1 | | E | E | E | 1-deoxy-D-xylulose-5-phosphate synthase |
| Cz05g23010.t1 | | E | E | E | isopentenyl-diphosphate synthase and dimethylallyl-diphosphate synthase |
| Cz10g18010.t1 | | E | E | E | Acetolactate enzymes |
| Cz17g13100.t1 | RBCS | E | E | NE | ribulose-bisphosphate carboxylase |
| Cz04g15140.t1 | | E | E | E | orotidine-5'-phosphate enzymes |
| Cz06g17080.t1 | | E | E | E | GMP enzymes |
| Cz09g30220.t1 | ACP@ | E | E | E | Lipid synthesis enzymes |
| Cz03g17210.t1 | | E | E | E | Folate reductases |
| Cz05g25150.t1 | | E | E | E | ribokinases |
| Cz10g09280.t1 | | E | E | E | ribosyltransferases |
| Cz08g13040.t1 | | E | E | E | glutamate-cysteine ligase |
| Cz15g21040.t1 | | E | E | E | glutathione synthase |
| Cz01g13260.t1 | BTA | E | E | E | betaine lipid synthase |
| Cz16g02090.t1 | | E | E | E | Lipid 3-phosphate o-acyltransferases |
| Cz09g31330.t1 | GPAT2 | E | E | E | acyl-coa acyltransferase |
| Cz11g03260.t1 | GPAT1 | E | E | E | glycerol-3-phosphate acyltransferase |
| Cz08g30040.t1 | MGD | E | E | E | galacto-glycerol enzymes |
| Cz07g23140.t1 | SQD2 | E | E | E | sulfolipid enzymes |
| Cz03g31030.t1 | SQD1 | E | E | E | UDP-sulfoquinovose synthase |
| Cz17g13240.t1 | PIS | E | E | E | CDP-diacylglycerol: myo-inositol 3-phosphatidyltransferase |
| Cz11g15030.t1 | ETK | E | E | E | choline kinase |
| Cz05g09130.t1 | EPT | E | E | E | Lipid phosphotransferases |
| Cz01g30230.t1 | | E | E | E | phospholipase |
| Cz06g24140.t1 | SDC | E | E | E | histidine and serine decarboxylase |
| Cz04g37140.t1 | | E | E | E | homoserine kinase |
| Cz17g17150.t1 | | E | E | E | threonine synthase |
| Cz04g20200.t1 | | E | E | E | aspartate-semialdehyde dehydrogenase |
| Cz09g04070.t1 | | E | E | E | 5-Amino-2-oxopentanoate:2-oxoglutarate aminotransferase |
| Cz04g39090.t1 | | E | E | E | 1-(5-phospho-D-ribosyl)-ATP:diphosphate phospho-alpha-D-ribosyl-transferase |

| | | | | | |
|---|---|---|---|---|---|
| Cz01g18040.t1 | | E | E | E | Histidinol enzymes |
| Cz10g04180.t1 | | E | E | E | imidazoleglycerol-phosphate dehydratase |
| Cz14g11090.t1 | | E | E | E | phosphoribosyl-AMP ezymes |
| Cz02g26110.t1 | | E | E | E | 1-(5-phosphoribosyl)-5-[(5-phosphoribosylamino)methylideneamino]imidazole-4-carboxamide isomerase, |
| Cz04g11190.t1 | | E | E | E | LL-diaminopimelate aminotransferase |
| Cz10g07220.t1 | | E | E | E | hydrodipicolinate synthases |
| Cz01g01020.t1 | | E | E | E | adenosylhomocysteinase |
| Cz07g08110.t1 | | E | E | E | acireductone dioxygenase |
| Cz07g16110.t1 | | E | E | E | acireductone synthase |
| Cz10g16160.t1 | | E | E | E | ATP:nicotinamide-nucleotide adenylyltransferases |
| Cz15g10170.t1 | | E | E | E | quinolinate synthase |
| Cz08g19150.t1 | | E | E | E | nicotinate-nucleotide diphosphorylase (carboxylating) |
| Cz04g33180.t1 | | E | E | E | 5-methyltetrahydrofolate:NAD+ oxidoreductase |
| CzMTg00230.t1 | | NE | NE | E | F0F1-ATP synthase Complex V |
| Cz06g10150.t1 | | NE | NE | E | F0F1-ATP synthase Complex V |
| Cz06g37200.t1 | | NE | NE | E | F0F1-ATP synthase Complex V |
| Cz18g09050.t1 | | NE | NE | E | F0F1-ATP synthase Complex V |
| Cz04g09240.t1 | | NE | NE | E | F0F1-ATP synthase Complex V |
| Cz03g14200.t1 | OSCP | NE | NE | E | F0F1-ATP synthase Complex V |
| CzMTg00140.t1 | | NE | NE | E | F0F1-ATP synthase Complex V |
| Cz18g13100.t1 | | NE | NE | E | F0F1-ATP synthase Complex V |
| UNPLg00020.t1 | | NE | NE | E | F0F1-ATP synthase Complex V |
| Cz06g20230.t1 | | NE | NE | E | F0F1-ATP synthase Complex V |
| Cz03g10280.t1 | CGLD22 | NE | NE | E | F0F1-ATP synthase Complex V |
| Cz05g29070.t1 | | NE | NE | E | F0F1-ATP synthase Complex V |
| Cz13g06290.t1 | | E | E | E | 2,3-Dihydroxy-3-methylbutanoate hydrolayses |
| Cz03g04080.t1 | TRK | E | E | E | transketolases |
| Cz14g05060.t1 | | E | E | E | Chorismate mutase |
| Cz04g15280.t1 | AROE | E | E | E | Shikimate enzymes |
| Cz13g16180.t1 | | E | E | E | phosphoribosyltransferase |
| Cz04g33170.t1 | | E | E | E | Chorismate synthase |
| Cz01g22210.t1 | | E | E | E | Indole derivatives enzymes |
| Cz05g19120.t1 | | E | E | E | Indole derivatives enzymes |
| Cz05g17190.t1 | | E | E | E | indole-3-glycerol-phosphate synthase |
| Cz13g09120.t1 | | E | E | E | Prephenate dehydratase and arogenate hydrolyase |
| Cz03g06230.t1 | DVR | E | E | E | divinyl chlorophyllide vinyl-reductase |
| Cz02g24270.t1 | CHLM | E | E | E | protoporphyrin IX methyltransferase |

| | | | | | |
|---|---|---|---|---|---|
| Cz08g28070.t1 | CHLG | E | E | E | chlorophyll synthase |
| Cz03g02090.t1 | GTR | E | E | E | glutamyl-tRNA reductase |
| Cz16g08310.t1 | CRD1 | E | E | E | Mg-protoporphyrin IX monomethyl ester cyclases |
| Cz09g03180.t1 | ALAD | E | E | E | porphobilinogen synthase |
| Cz02g42010.t1 | PPOX | E | E | E | protoporphyrinogen IX oxidase |
| Cz14g24240.t1 | UROS1 | E | E | E | uroporphyrinogen III synthase |
| Cz05g10280.t1 | | E | E | E | AMP-lyases |
| Cz10g16210.t1 | | E | E | E | phosphoribosylaminoimidazolecarboxamide formyltransferase and IMP cyclohydrolase |
| Cz05g31230.t1 | | E | E | E | phosphoribosylaminoimidazole carboxylase |
| Cz17g13050.t1 | | E | E | E | adenylosuccinate synthase |
| Cz10g19230.t1 | | E | E | E | oxidized-thioredoxin 2'-oxidoreductases |
| Cz16g14140.t1 | | E | E | E | phosphoribosylaminoimidazolesuccinocarboxamide synthase |
| Cz12g03220.t1 | | E | E | E | phosphoribosylformylglycinamidine synthase |
| Cz16g22110.t1 | | E | E | E | dihydroorotase |
| Cz01g01200.t1 | | E | E | E | Deoxy kinases |
| Cz14g10040.t1 | | E | E | E | UTP ligases |
| Cz15g07180.t1 | | E | E | E | 2-isopropylmalate synthase |
| Cz12g04120.t1 | | E | E | E | 5-amino-6-(5-phosphoribitylamino)uracil:NADP+ 1'-oxidoreductase |
| Cz06g12170.t1 | | E | E | E | riboflavin production |
| Cz08g11170.t1 | | E | E | E | 2-Methyl-4-amino-5-hydroxymethylpyrimidine-diphosphate enzymes |
| Cz12g09160.t1 | | E | E | E | Divalent cation (Mg2+) transport system |
